# PERK modulation, with GSK2606414, Sephin1 or salubrinal, failed to produce therapeutic benefits in the SOD1G93A mouse model of ALS

**DOI:** 10.1101/2023.09.17.558143

**Authors:** Fernando G. Vieira, Valerie R. Tassinari, Joshua D. Kidd, Andrew Moreno, Kenneth Thompson, Steven Perrin, Alan Gill, Theo Hatzipetros

## Abstract

Amyotrophic lateral sclerosis (ALS) has been linked to overactivity of the protein kinase RNA-like ER kinase (PERK) branch of the unfolded protein response (UPR) pathway, both in ALS patients and mouse models. However, attempts to pharmacologically modulate PERK for therapeutic benefit have yielded inconsistent and often conflicting results. This study sought to address these discrepancies by comprehensively evaluating three commonly used PERK modulators (GSK2606414, salubrinal, and Sephin1) in the same experimental models, with the goal of assessing the viability of targeting the PERK pathway as a therapeutic strategy for ALS. To achieve this goal, a tunicamycin-challenge assay was developed using wild-type mice to monitor changes in liver UPR gene expression in response to PERK pathway modulation. Subsequently, multiple dosing regimens of each PERK modulator were tested in standardized, well-powered, gender-matched, and litter-matched survival efficacy studies using the SOD1G93A mouse model of ALS. The alpha-2-adrenergic receptor agonist clonidine was also tested to elucidate the results obtained from the Sephin1, and of the previously reported guanabenz studies, by comparing the effects of presence or absence of α-2 agonism. The results revealed that targeting PERK may not be an ideal approach for ALS treatment. Inhibiting PERK with GSK2606414 or activating it with salubrinal did not confer therapeutic benefits. While Sephin1 showed some promising therapeutic effects, it appears that these outcomes were mediated through PERK-independent mechanisms. Clonidine also produced some favorable therapeutic effects, which were unexpected and not linked to the UPR. In conclusion, this study highlights the challenges of pharmacologically targeting PERK for therapeutic purposes in the SOD1G93A mouse model and suggests that exploring other targets within, and outside, the UPR may be more promising avenues for ALS treatment.

## Introduction

Accumulations of misfolded, unfolded, or aggregated proteins can trigger endoplasmic reticulum (ER) stress, activating the unfolded protein response (UPR), a process known to be involved in the pathophysiology of neurodegenerative disorders like amyotrophic lateral sclerosis (ALS). The UPR is a tightly regulated signal transduction pathway, modulating many ER processes, the goal of which is to restore protein homeostasis by either an attenuation of translation, a recruitment of chaperones needed for protein folding or an initiation of degradation pathways if the ER stress persists^1^. The UPR is composed of three interconnected branches, each mediated by a different ER membrane stress sensor: the protein kinase RNA-like ER kinase (PERK) branch, the transmembrane basic leucine zipper activating transcription factor-6 (ATF6) branch and the inositol-requiring protein-1 branch (IRE1) shown in Fig 1.

**Fig 1:**
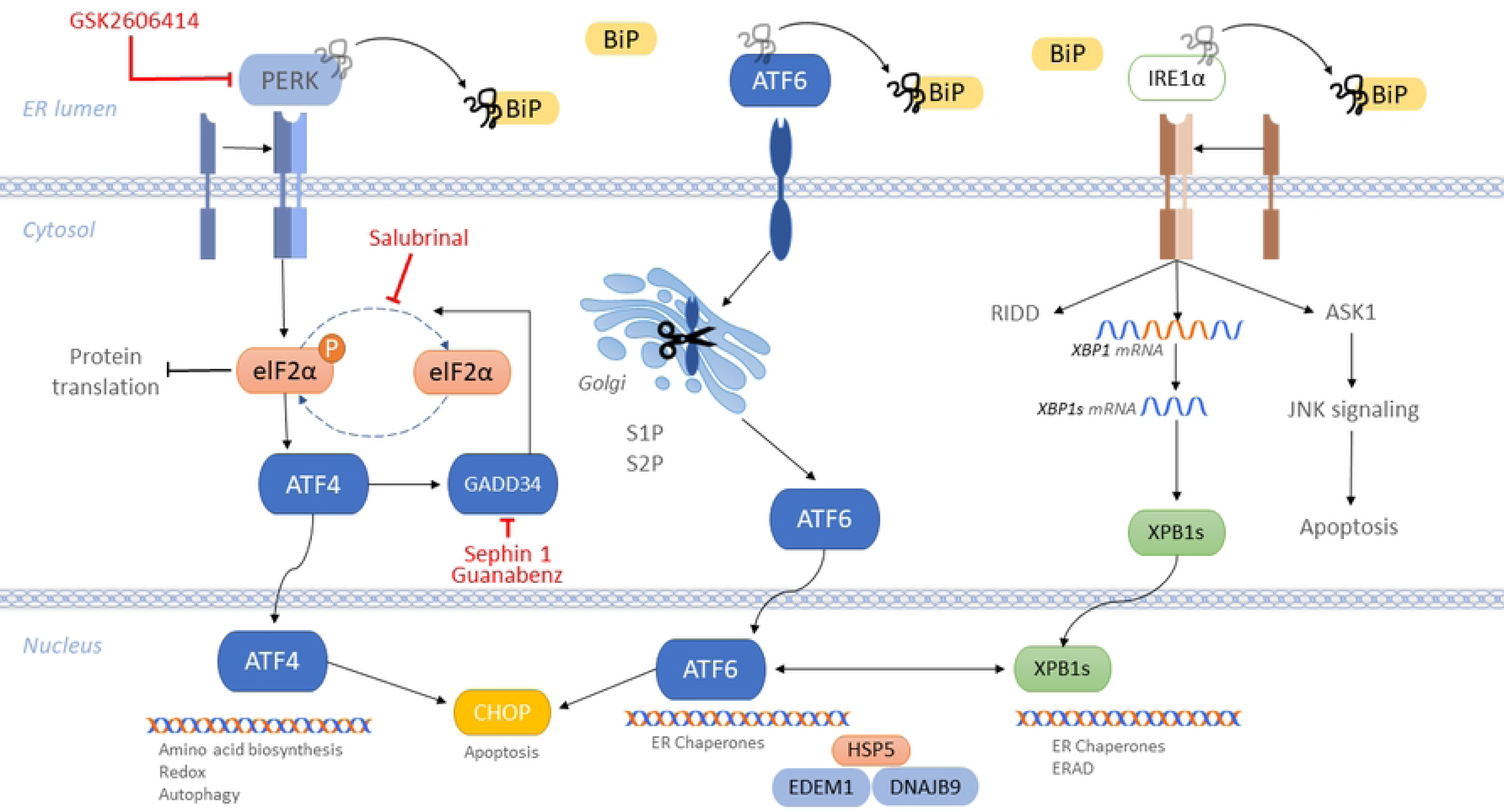
Illustration depicting the intricate mechanism of the unfolded protein response (UPR). The drugs discussed in this study, accompanied by their respective sites of action, are shown in red.

In response to ER stress, the chaperone binding immunoglobulin protein (BiP) preferentially binds the abundant aberrant proteins, physically dissociating itself from PERK, IRE1, and ATF6 and thereby permitting their activation. The *PERK branch* is the branch that elicits the most rapid response to stress, protecting the ER by reducing its folding load^2^. Once activated, PERK leads to the attenuation of general protein synthesis through a phosphorylation of eukaryotic translation initiation factor 2α (eIF2α) and the subsequent upregulation of selected stress-induced mRNAs, such as the transcription factor ATF4, which translocates into the nucleus and induces the expression of genes involved in amino acid biosynthesis, resistance to oxidative stress, and autophagy^3^. Growth arrest and DNA damage 34 (GADD34) dephosphorylates p-eIF2α, allowing protein synthesis to return to normal. However, when the UPR is unable to resolve the accumulation of aberrant proteins, the pro-apoptotic transcription factor C/EBP homologous protein (CHOP) is activated^4^. The *ATF6 branch* is activated when ATF6, upon its release from BiP, is directed to the cis-Golgi, where it is proteolytically cleaved becoming active. The active ATF6 fragment in turn activates the transcription of several other proteins including X-box binding protein 1 (XBP1), and CHOP. The *IRE1 branch* is the most evolutionary conserved branch of the UPR. There are two IRE1 isoforms, IRE1α and IRE1β, IRE1α being the predominant isoform in the UPR. Active IRE1α drives the splicing of a 26-nucleotide long section of mRNA from XBP1, which restores the translation of XBP1 (XBP1s)^3^. This enhances the expression of UPR-dependent genes, including a gene for ER folding (ERAD), ER degradation-enhancing alpha-mannosidase-like protein 1 (EDEM1), and others^5^. Through XBP1-independent pathways, IRE1 binds to tumor necrosis factor (TNF) receptor-associated factor 2 (TRAF2) and induces JUN amino-terminal kinase (JNK) activation, via ASK1. This interaction is known to modulate autophagic and apoptotic cell death^6^.

A preponderance of evidence suggests that the three UPR branches are overactive in ALS patients, albeit most of the evidence comes from post-mortem samples^7,8,9^. Spinal cords of sporadic ALS patients were found to have significant upregulation of IRE1, PERK and ATF6 compared to controls without neurological disease^10^, as well as increased expressions of ATF4 and XBP-1, which are downstream from PERK and IRE1 respectively^11,12^. Moreover, CHOP was increased in the spinal cord tissue of ALS patients compared to controls^13^. More recently, a study looking at the peripheral blood mononuclear cells of patients living with sporadic ALS found a significant upregulation of spliced XBP1, and of the two stress sensors IRE1α and ATF6, compared to healthy controls, indicating that UPR induction is not a terminal or near-terminal event but rather a compensatory mechanism, that begins to happen early in the course of the disease^14^. Approximately half of the patients in this study were within a year of ALS diagnosis, and the average disease duration was twenty months. Notably, in the same study the levels of PERK were not significantly increased.

Studies in animal models have further reinforced the link between UPR induction and ALS^15,16,17^. Specifically, in different strains of mutant SOD1 transgenic mice, activation of IRE1α, CHOP and XBP-1 mRNA splicing have all been repeatedly observed^18,19,20^. Genetic studies in ALS mouse models, primarily targeting the PERK pathway, showed that UPR gene manipulation can have a great impact on the progression of the disease, though sometimes with contrasting results. In one such study, PERK haploinsufficient mice were crossed with transgenic SOD1-G85R mice and the progeny displayed accelerated disease, suggesting that diminished capacity to respond to ER stress, via PERK activation, is deleterious^21^. In a subsequent study, SOD1 transgenic mice harboring a GADD34 mutation were developed, with a decreased capacity for dephosphorylation of the eIF2 alpha subunit, and they had a delayed and less severe disease phenotype^22^. Knocking down GADD34, by administering AAV9-encoded GADD34 shRNA into neonatal, but not adult, SOD1G93A or SOD1G85R mice also led to a significantly increased survival^23^, providing more support for the PERK pathway’s involvement in the pathology of SOD1 mouse models. In contrast, a more recent genetic study, involving five different lines of transgenic mice expressing human SOD1 mutations, demonstrated that PERK haploinsufficiency had no effect on the disease progression of mutant SOD1 mice. This suggests that the PERK pathway is not an ideal therapeutic target for mutated SOD1-induced ALS^24^.

The genetic validation of therapeutic targets within the UPR, stimulated interest in the development of small molecule therapeutics that selectively modulated UPR signaling, particularly the PERK branch. Drug discovery strategies often involved identifying promising PERK modulators through primary drug screening assays that model certain aspects of ER stress and UPR upregulation, followed by validation in animal models of protein-misfolding diseases. ER stressors like hypoxia, glucose deprivation and certain chemicals like tunicamycin (TUN), all induce UPR upregulation in a host of mammalian and other cell types and thus these *in vitro* models have been widely used as screening assays for identifying UPR homeostasis-restoring compounds^25^. This approach has identified several small molecules, and while some preclinical studies have demonstrated promising results, the outcomes have frequently been inconsistent and even conflicting. What is intriguing is that compounds with opposing mechanisms of action have occasionally yielded similar outcomes, and conversely, compounds with similar mechanisms of action have produced different results.

Salubrinal was one of the first PERK pathway modulating small molecules that showed promise as an ALS therapeutic. Salubrinal was shown to mitigate ER stress by delaying the dephosphorylation of eIF2α and enhancing the activity of PERK (site of action indicated in Fig 1). It was shown to suppress the formation of insoluble SOD1 aggregates, to attenuate the cytotoxicity in cells overexpressing SOD1*in vitro*^26^, and to attenuate the disease symptoms and prolong the survival of SOD1G93A mice^27^. However, despite its initial promising results, subsequent studies in SOD1 mouse models have not been able to replicate the beneficial effects of salubrinal. The reasons for this discrepancy are not clear but may be due to the unfavorable pharmacological profile of the drug or simply due to unpublished negative results in subsequent studies. Regardless, further exploration is needed to reach a consensus on the potential of salubrinal as an ALS treatment.

GSK2606414, also known as compound 38, was the first selective PERK inhibitor to be discovered. Chronic ER stress, as seen in ALS, promotes pro-apoptotic signaling downstream of PERK and thus the blockade of PERK signaling was anticipated to result in neuroprotection. GSK2606414, in contrast to salubrinal, blocked PERK activation by targeting its kinase active site and inhibiting its autophosphorylation step, which is required for activation (site of action indicated in Fig 1). The compound demonstrated good oral bioavailability and blood-brain barrier penetration and showed promising results as a neuroprotective agent in a fly model of TDP-43 proteinopathy^28^ and a prion disease mouse model^29^. However, despite its promising results, GSK2606414’s development as a therapeutic was unfortunately halted due to its systemic effects, caused by an inhibition of PERK in the pancreas, resulting in slight weight loss and increased blood sugar levels, though they remained below diabetic levels in mice. Despite encountering setbacks in its development, GSK2606414 may still hold promise as a therapeutic for ALS since the potential benefits that it could bring to ALS patients outweigh what may be relatively mild risks.

Guanabenz, an alpha-2 adrenergic receptor agonist and an FDA-approved antihypertensive, was shown to interact with a regulatory subunit of the protein phosphatase GADD34 (site of action indicated in Fig 1), selectively disrupting the dephosphorylation of the alpha subunit of eIF2α^30^ and was initially demonstrated to be efficacious in the SOD1G93A mouse model of ALS^31,32^. However, a later study contradicted these reports, showing not just a lack of benefit, but an accelerated ALS-like disease progression in a strain of mutant SOD1 transgenic mice^33^. This discrepancy was attributed to the counteracting actions of guanabenz on the alpha-2 adrenergic receptor, which negated its UPR modulating effects. Moreover, a recent phase 2 clinical trial with guanabenz in ALS patients, produced underwhelming evidence in support of further clinical studies and instead indicated that a larger trial with a different UPR-targeting molecule, free of the alpha-2 adrenergic side-effect profile of guanabenz, was warranted^34^.

Sephin1 is a close analog of guanabenz that lacks its alpha-2 adrenergic receptor affinity. Like guanabenz, Sephin1 is also a selective inhibitor of the phosphatase regulatory subunit GADD34, prolonging the benefit of eIF2α phosphorylation, which in essence activates the PERK pathway (site of action indicated in Fig 1). Chronic administration of Sephin1 showed protective effects in Charcot-Marie-Tooth disease and in the SOD1G93A mouse model of ALS^35^. Sephin1 also delayed the onset of clinical symptoms in a multiple sclerosis mouse model^36^, and extended the survival of prion infected mice^37^. Sephin1’s neuroprotective potential has not yet been questioned; however recent doubts have arisen regarding its mechanism of action and its proposed targets, which still need further investigation^38,39^.

Despite numerous genetic studies validating the involvement of UPR in ALS, no small molecule therapeutics targeting the UPR have been developed as drugs for ALS thus far. One reason for this might be that the primary screening assays used to identify these compounds were not adequately predictive of *in vivo* efficacy. None of these assays perfectly capture all aspects of UPR biology, and as a result, “hits” do not always repeat, even across similar assays. Another reason might be the suboptimal design and interpretation of the preclinical studies used to validate UPR-modulators. It has been well documented that flaws in preclinical animal research design are a recurring problem, especially in the context of ALS mouse models^40^. Deciphering the reasons for the inconsistent outcomes of PERK modulators, based on existing data, is nearly impossible since each molecule was tested in different models under different conditions.

The aim of the current study was to reconcile the inconsistent outcomes of promising PERK modulators, namely GSK2606414, salubrinal, and Sephin1, by conducting side-by-side testing in ALS-relevant assay systems, and ultimately determining whether PERK is a promising pharmacological target for ALS. To determine whether the lack of translatability of *in vitro* screening assays was contributing to some of the inconsistent *in vivo* outcomes, we developed a mouse-based tunicamycin assay that we hypothesized would better predict *in vivo* efficacy. To determine whether an improved preclinical study design would confirm and validate the beneficial effect of PERK modulators, as seen by others, we conducted a large-scale effort comprised of eight survival efficacy studies in SOD1G93A mice. These were performed according to internationally established quality standards^41,42^ assessing the therapeutic potential of GSK2606414, salubrinal, and Sephin1. Finally, we sought to probe whether the lack of therapeutic effect that we previously demonstrated for guanabenz in SOD1G93A mice could be related to its alpha-2-adrenergic receptor agonist properties. To do this we performed a survival efficacy study with the alpha-2-adrenergic agonist clonidine, hypothesizing that it would accelerate disease progression in this model.

## Materials and Methods

### Animals

The SOD1G93A mouse colony that provided mice for the current study was derived from the high copy B6SJLTgN (SOD1G93A)1Gur strain originally produced by Gurney, *et al*.^43^, and obtained from The Jackson Laboratory (Bar Harbor, Maine). The colony was maintained by Biomere (Worcester, MA) by crossing hemizygous C57BL6-SJL sires harboring the SOD1 transgene with non-transgenic C57BL6-SJL dams. SOD1G93A mice and wild-type litter mates were shipped to ALS TDI at 35–45 days of age, then genotyped for the expression of the transgene as previously described^44^, and allowed at least one week to acclimate to ALS TDI’s animal facility (12-h light/dark cycle at a temperature of 18–23 °C and 40–60% humidity) before being assigned to a study. Male mice were singly housed, because of aggressive behavior, while female mice were housed in pairs. Environmental enrichment was provided in the form of plastic huts. Food and water were provided *ad libitum* and the standard diet used was Teklad Global diet #2918 for rodents (Harlan Laboratories, Houston, TX). Whenever a treatment was formulated in chow, female mice were also singly housed so that the food intake of each mouse could be determined. All the procedures described in this research were performed by Assistant Laboratory Animal Technician (ALAT)-certified personnel in a facility that was regularly inspected by a consultant veterinarian. All the studies were approved by the ALS TDI Institutional Animal Care and Use Committee (IACUC) and in accordance with the Institute for Laboratory Animal Research (ILAR) Guide for Care and Use of Laboratory Animals^45^.

### Drugs and Chemicals

Tunicamycin (TUN), N-methyl-2-pyrrolidone (NMP), Kolliphor® HS15, methylcellulose and clonidine hydrochloride were purchased from Sigma-Aldrich (St. Louis, MO). Salubrinal was purchased from Tocris, now part of Bio-Techne Corporation (Minneapolis, MN). Sephin1 acetate was purchased from Otava chemicals (Concord, ON, Canada). GSK2606414 was synthesized by Perfemiker Canspec China (Shanghai, China). Refer to S1 Fig for compound structures of cited compounds.

### Gene expression analyses

#### In wild-type mice, in the context of the *in vivo* TUN assay

RNA was extracted from wild-type mouse livers using the Agencourt RNAdvance Tissue kit (Beckman Coulter, Brea, CA), and cDNA was synthesized using the High-Capacity cDNA Reverse Transcription Kit (Applied Biosystems, Waltham, MA). RT-qPCR was performed using TaqMan low density array (LDA) cards. First, reference gene transcripts were selected as internal controls. Liver RNA samples, from TUN-treated and untreated mice, were used in an RT-qPCR experiment on the TaqMan Mouse Endogenous Control Array (Thermo Fisher Scientific, Waltham, MA). The resulting Ct’s were analyzed using GraphPad Prism to find the coefficient of variation of each gene across the samples and to ascertain how well each gene predicted template concentration. Gusb, 18S, Hprt, Polr2a, and Tfrc were determined to be the genes with the least variance and therefore were selected as “control genes”. Then, custom LDA cards were designed which included the five “control genes” as well as the following “target genes”, in duplicate: CHOP, HSPA5, GADD34, DNAJB9, ATF6, ATF4, PERK, EDEM1, XBP1, IRE1α. All gene expression analyses were performed using the custom LDA cards. Resulting Ct values were normalized to relative expression values using the geNorm software as previously described^46^.

#### In SOD1G93A mice of different ages

Spinal cords were collected from groups of SOD1G93A animals and wild-type littermates at 30, 50, 60, 80, 90, 100, 110 and 120 days of age. For each time point, five non-transgenic and five SOD1G93A tissues were collected and processed independently, for a total of 70 tissues samples. Genome-wide transcriptional profiling using Affymetrix MOE430vII gene chips was performed. For this research we focused on UPR gene expression.

### Drug Survival Efficacy Studies

Eight different survival efficacy studies, in SOD1G93A mice, were performed for this project. Each of these studies contained 64 mice which were divided into either a drug-treated (n=32) or a vehicle-treated (n=32) group. The two groups were age-matched, litter-matched, and gender-balanced. For all studies, hSOD1G93A transgene copy number was verified and only mice within an acceptable gene copy range, known to produce the disease phenotype, were included. For all survival efficacy studies, drug treatments began at around age 50±3 days and lasted for approximately 70-80 days until the “humane endpoint” was reached, which is defined as the time when the mice were unable to right themselves when placed on their sides for 10 seconds. At this stage the mice were unable to feed themselves and they were immediately euthanized. This usually occurred at age 123±5 days in males and at age 127±5 days in females. Among the 512 mice included in our studies, 504 succumbed to the progression of their disease upon reaching the humane endpoint. Six mice were intentionally euthanized before reaching this endpoint due to injuries stemming from oral gavage administration errors, while an additional two mice were sacrificed earlier than the humane endpoint due to unresolved tail infections following consultation with our veterinarian. Euthanasia was performed, in a different room than their holding room, according to the guidelines of The American Veterinary Medical Association (AVMA), by using a gas chamber with 100% CO_2_, at a flow rate of approximately 20% of the chamber volume per minute.

Mouse behavior and overall health was monitored once daily during the pre-symptomatic phase of their disease (from approximately age day 50 to age day 110) and twice daily during the symptomatic phase of their disease (from approximately age day 110 to the humane endpoint). With the exception of the eight mice mentioned earlier, which experienced oral gavage injuries and tail infections, no indications of pain or discomfort were detected in the remaining mice. Body weights and neurological scores (NeuroScores or NS) were assessed daily, at about the same time, on a bench-top in the animal holding room. Body weights were measured using a scale with an accuracy of one tenth of a gram. NeuroScores were determined, as in a previously described protocol^47^, by observing the hindlimbs of mice while being suspended by the tail, while walking, or when placed on their side.

Briefly, NS were assigned for each hindlimb (left or right) independently on a scale from 0 to 4 corresponding to the following set of observations: NS 0 indicated that the hindlimb presented a normal splay (Pre-symptomatic); NS 1 indicated the appearance of an abnormal hindlimb splay but that the gait was not severely affected (First symptoms); NS 2 indicated that the hindlimb was partially or completely collapsed and even though the hindlimbs were used for forward motion there was foot-dragging or toe-curling observed (Onset of paresis); NS 3 indicated that there was rigid paralysis in the hindlimb and it was not being used for forward motion (Paralysis); NS 4 indicated that there was rigid paralysis in the hindlimbs and that the mouse was unable to right itself up within 10 seconds when placed on its side (Humane end-point).

All body weight and NeuroScore data captured during the study were entered into a custom ALS TDI Laboratory Information Management System (LIMS). These data were used to compute the main four readouts of a survival efficacy study: body weight maintenance, neurological disease progression, disease onset and survival duration.

### Drug Formulations

Test compounds were formulated and administered to mice as follows:

#### Salubrinal

Two drug efficacy studies with salubrinal were performed. In the first study salubrinal was formulated in 2.5% NMP, 2.5% Kolliphor HS15/95% filtered tap water at a concentration of 0.5 mg/mL or 5 mg/kg. In the second study, salubrinal was formulated in 5% NMP/ 5% Kolliphor HS15 / 90% filtered tap water at a concentration of 1.5 mg/mL or 15 mg/kg. The vehicle treatments were 2.5% NMP/ 2.5% Kolliphor HS15/95% filtered tap water and 5% NMP/5% Kolliphor HS15/90% filtered tap water respectively. In both studies salubrinal was administered intraperitoneally, once daily, at a volume of 10 mL/kg body weight.

#### GSK2606414

Three drug efficacy studies with GSK2606414 were performed. In two of them GSK2606414 was suspended in 1% methylcellulose and administered via oral gavage, once daily, at a concentration of either 1.8 mg/mL or 5 mg/mL, corresponding to 18 mg/kg or 50 mg/kg when adjusted for body weight dosing. Injection volumes for these studies were 10 mL/kg body weight. In the third study, GSK2606414 was added to chow (18% protein Teklad global diet) with the goal of delivering 50 mg/kg/day, based on a 4 g average daily *ad libitum* chow consumption. The vehicle-treated group received a standard 18% protein Teklad global diet.

#### Sephin1 acetate

Two drug efficacy studies with Sephin1 acetate were performed. In the first study, Sephin1 acetate was formulated in 5% glucose dissolved in water to a concentration of 0.4 mg/mL, corresponding to 4 mg/kg when adjusted for body weight dosing, and was administered intraperitoneally every other day. In the second study, Sephin1 acetate was formulated in 0.9% sterile physiological saline to a concentration of 1 mg/mL, corresponding to 10 mg/kg when adjusted for body weight dosing, and was administered intraperitoneally once daily. The vehicle groups were treated with 5% glucose dissolved in water and 0.9% sterile physiological saline, respectively. Intraperitoneal injection volumes for these studies were 10 mL/kg body weight.

#### Clonidine hydrochloride

Clonidine was formulated in 0.9% sterile physiological saline to a concentration of 0.05 mg/mL and administered intraperitoneally every other day. The vehicle treatment group received 0.9% sterile physiological saline. Injection volumes for both clonidine hydrochloride and vehicle-treated cohorts were 10 mL/kg body weight.

### Statistical Analyses

#### Gene Expression data

Changes in gene expression after tunicamycin challenge were expressed as fold-change of saline-treated control animals. Changes in gene expression, as a result of drug pretreatment, were compared using Dunnett’s test for multiple comparisons to a control group. Data were also tested for dose-level difference using the Tukey HSD test for all-pair comparisons.

#### Disease Progression over Time, Effect of Treatment on Neurological Severity Score

Daily neurological severity scores were taken from study start until death. These ordinal scores ranging from 0 to 4 were modeled in relation to the animal’s median age at each score level using ordinal logistic regression. The model fits cumulative response probabilities to the logistic function of a linear model using maximum likelihood. Likelihood-ratio and Wald Chi-square test probabilities are provided for the treatment effect.

#### Analyses Time to Event Data for Disease Onset and Survival Duration

Time to onset of definitive disease and survival time were analyzed using the Kaplan-Meier and Cox proportional hazard methods. Kaplan-Meier survival fit analysis employed the Log-Rank and Wilcoxon tests for statistical significance. Cox proportional hazards analysis was also performed to determine hazard ratios and test for statistical significance of their differences using the Effect Likelihood Chi Square test.

Statistical analyses were performed using JMP 17.1.0, SAS Institute, Inc., SAS Campus Drive, Cary, NC 27513, USA. Cox proportional hazard fitting, using litter as a frailty term, was performed using Stata BE 18.0, 4905 Lakeway Drive, College Station, TX 77845, USA. p-values less than 0.05 were taken to be statistically significant.

## Results

### *In vivo* Tunicamycin Assay Development

To induce widespread activation of the UPR, we used the antibiotic tunicamycin (TUN), which blocks N-glycosylation and creates ER stress. TUN (1 mg/kg, IP) or saline were administered to wild-type mice and six hours post-administration the relative expression of UPR genes was measured in the liver. This specific TUN regimen was chosen after conducting preliminary dose-response studies (data not shown). The liver tissue was used as a surrogate for the spinal cord or brain, which are the tissues of interest in ALS, because TUN does not readily cross the blood-brain-barrier and thus would not activate the UPR in the central nervous system.

A single dose of TUN led to a prevalent upregulation of UPR genes (Fig 2). The mRNA levels of genes from all three UPR branches were elevated in TUN-treated mice compared to those treated with saline. The ER membrane-resident sensors PERK, ATF6, and IRE1α showed a 6-, 8-, and 3-fold increase respectively. Meanwhile, ATF4 and GADD34, which are downstream from PERK, were upregulated 6- and 20-fold respectively, and CHOP, which is further downstream, was elevated 363-fold compared to saline-treated mice. HSPA5 and Dnajb9, members of Heat Shock Protein Families 70 (Hsp70) and 40 (Hsp40), were robustly upregulated 26-fold and 17-fold, respectively, compared to saline-treated mice. Edem1, associated with the activation of the ER-associated protein degradation (ERAD) system in response to ER stress, was also upregulated 4-fold by TUN. Finally, total mRNA levels of X-box-binding protein 1 (XBP1), a splicing target of IRE1α, were upregulated 4-fold.

**Fig 2:**
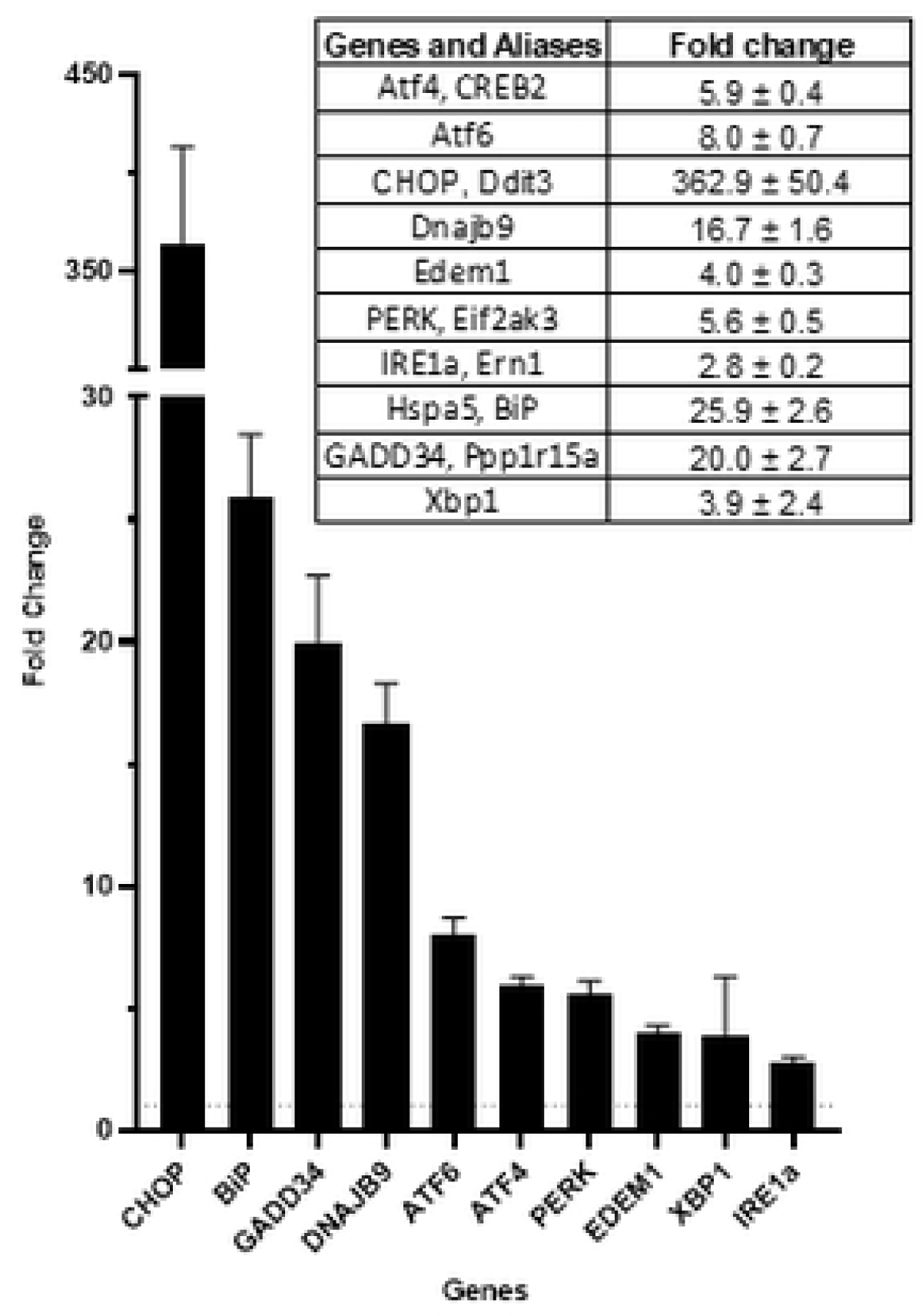
Impact of tunicamycin challenge (1mg/kg, IP) on the relative expression of UPR genes in the livers of WT mice. Fold change values depict gene expression in tunicamycin-treated mice relative to saline-treated mice, six hours after the treatments. The dataset comprises n=23 tunicamycin-treated and n=24 saline-treated mice. The averages are presented as mean ± SEM. The dotted horizontal line at y=1 represents the baseline expression level with no change in gene expression.

Notably, the gene expression changes caused by TUN in the livers of WT mice resembled the natural gene expression changes observed in the spinal cords of SOD1G93A mice because of their age-related disease progression (Fig 3). Although the TUN mouse-based assay is an artificial system with exaggerated effects, the directionality of the gene expression changes was similar in both TUN-treated mice and older SOD1G93A mice compared to their respective controls. Most UPR genes, such as ATF4, ATF6, CHOP (Ddit3), GADD34 (Ppp1r15a), EDEM1, and HSPA5 were upregulated in both TUN-treated mice and older SOD1G93A mice. Expression levels of the ER membrane-resident sensors PERK (eif2ak3) and IRE1α were not substantially altered in SOD1G93A mice, compared to age-matched WT controls, despite downstream genes in both branches being altered.

**Fig 3:**
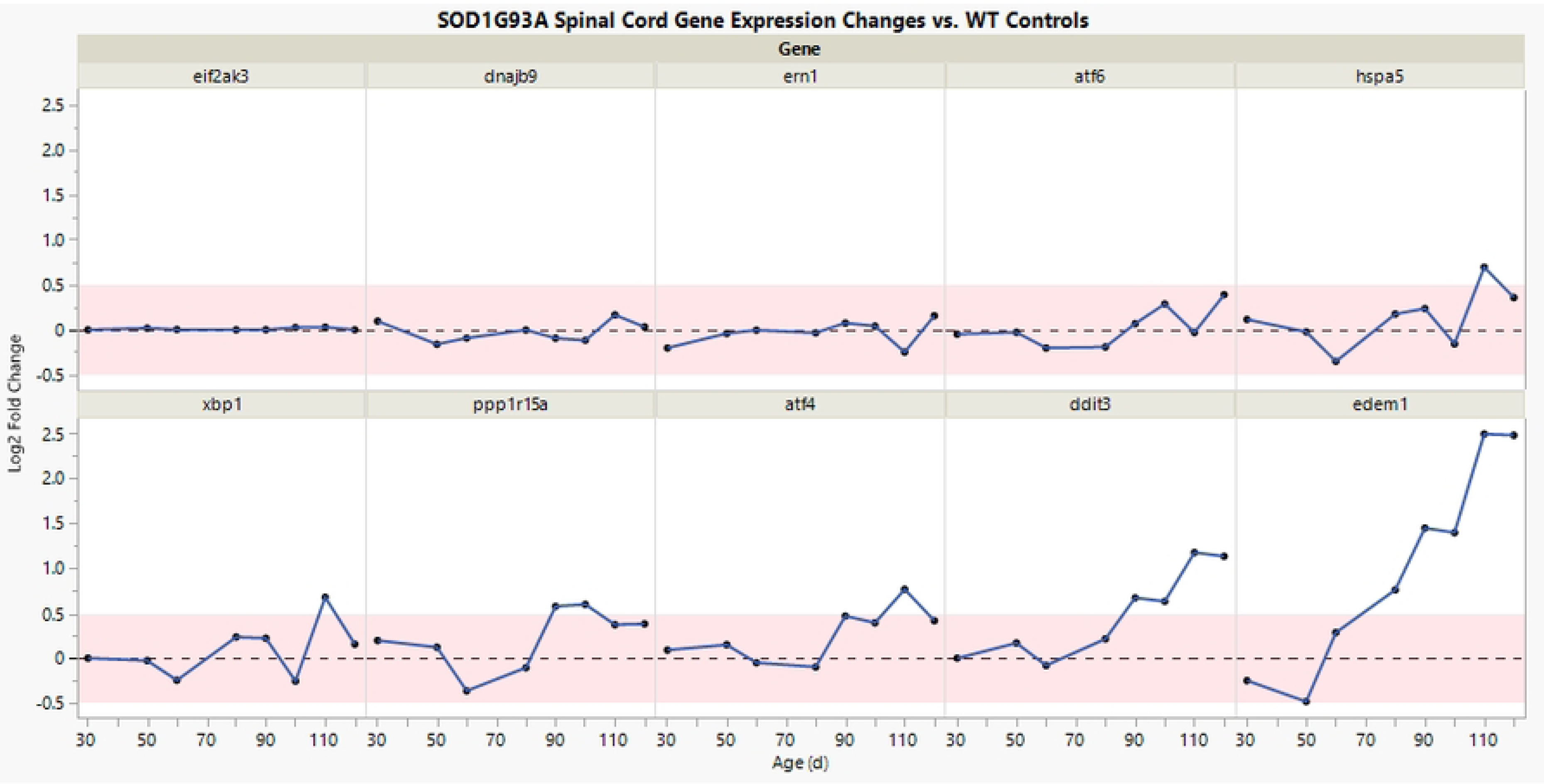
A time course of UPR gene expression in the spinal cord of SOD1G93A mice. Plot shows fold differences in gene expression (y-axis), comparing SOD1G93A mice to age-matched wild-type mice, as a function of their age in days (x-axis). Each UPR gene is depicted in a separate panel. The data shown are log2 fold change averages, based on n=5 SOD1G93A and n=5 WT mice, for each timepoint.

### Drug Screening in the *In vivo* Tunicamycin assay

Next, the effectiveness of GSK2606414, Sephin1 and salubrinal, in modulating the TUN-induced changes in gene expression was evaluated in the same assay. The UPR activity of these three compounds, thought to be mediated through its PERK branch, was previously described in *in vitro* assays^48,49^. Wild-type mice were treated with either a test compound or saline two hours before being subjected to the TUN challenge. TUN was administered at a dose of 1 mg/kg, IP, and 6 hours afterwards the mice were humanely euthanized, and their livers were collected for UPR gene expression analysis.

Of the compounds tested, the PERK inhibitor GSK2606414 was the one that most definitively blocked the effects of the TUN challenge, in a concentration dependent manner (Fig 4). Administered orally at 18 or 50 mg/kg, it reduced the overexpression of PERK branch genes ATF4 (by 23% and 45% at low and high dose, respectively), GADD34 (by 59% at high dose), and CHOP (by 42% and 70% at low and high dose, respectively). Additionally, high-dose GSK2606414 reduced overexpression of ATF6 (by 39%) and Hsp5 and Dnajb9 (by 44% and 41%, respectively). However, GSK2606414 did not affect the overexpression of genes in the IRE1α branch, including IRE1α and Xbp1.

**Fig 4.**
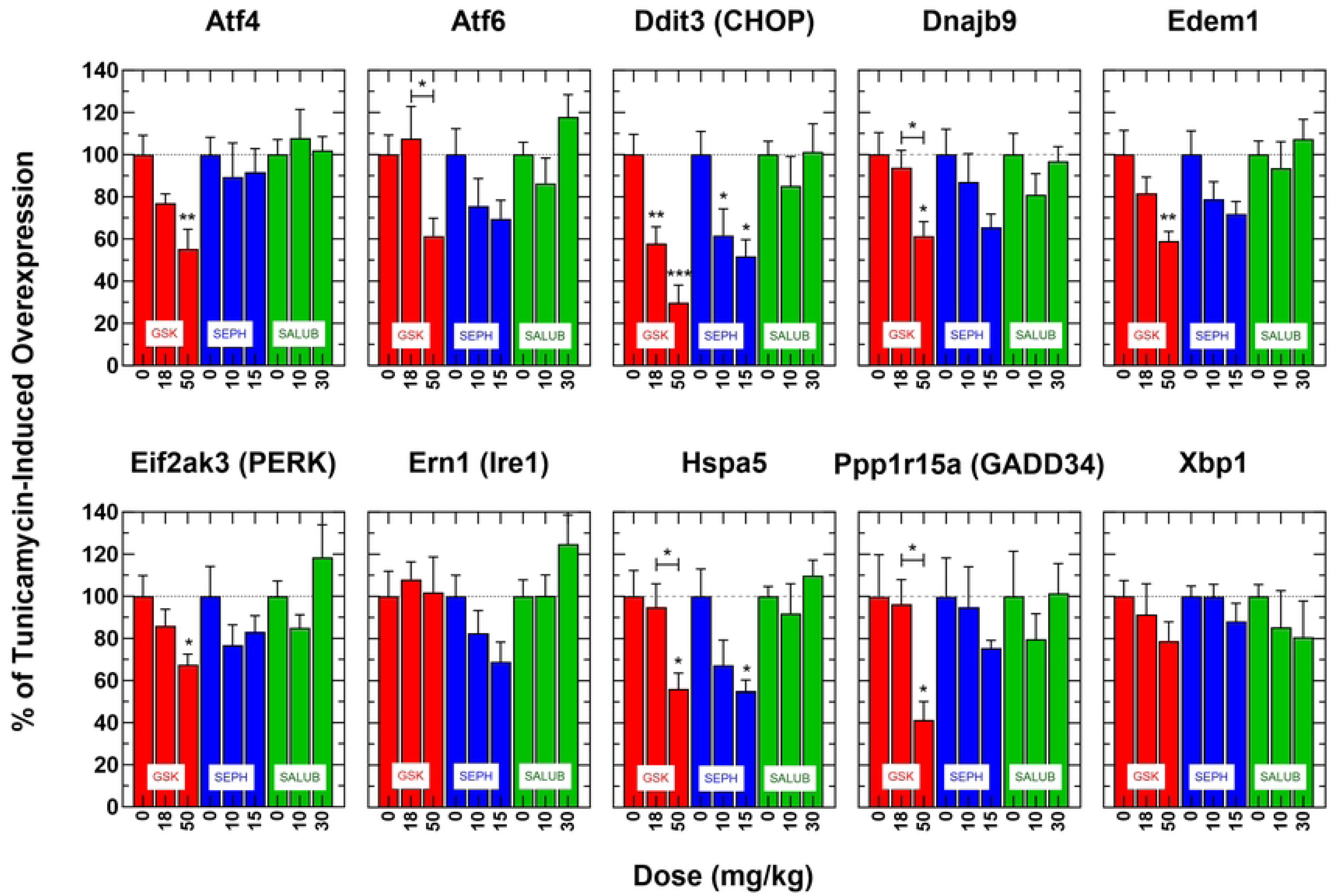
Effect of PERK modulators on liver UPR gene expression in mice challenged with tunicamycin. WT mice were pretreated with varying doses of GSK2606414, Sephin1, and salubrinal two hours prior to the tunicamycin challenge. The expression of each UPR gene is depicted in a separate panel. Each bar represents the percent of tunicamycin-induced overexpression of the gene measured without drug pretreatment (dose=0) or when pretreated at different dose levels of the tested drugs for each specific gene. Statistical analysis was performed using Dunnett’s test to compare data with the untreated control group (dose=0) and the Tukey HSD test to evaluate dose-dependent differences among all pairs. Significance levels: *=p<0.05, **=p<0.01, ***=p<0.001.

Sephin1 also attenuated some elements of the TUN challenge when administered intraperitoneally at 10 or 15 mg/kg. The high dose of 15 mg/kg significantly reduced the over-expression of several UPR genes, including Edem1 by 28%, Dnajb9 by 34%, Hsp5 by 45% and CHOP by 48%. Additionally, Sephin1 decreased the expression of IRE1α by 31% which was almost statistically significant (p=0.0946). Unlike GSK2606414, Sephin1 did not attenuate the expression of PERK branch genes. Out of all the compounds tested, Sephin1 was the only one that had any impact on the expression of the IRE1α branch (Fig 4).

Salubrinal did not have an effect on the expression of UPR genes. This was true for both doses tested, 10 and 30 mg/kg, given by intraperitoneal injection (Fig 4).

### GSK2606414 Survival Efficacy Studies

Given its activity in the *in vivo* tunicamycin assay, positive pharmacological properties, and supporting published data, we extensively tested GSK2606414 in three standardized drug efficacy studies in SOD1G93A mice.

GSK2606414 was initially evaluated at a dose of 50 mg/kg through daily oral administration. This dose was chosen as it had previously shown efficacy in a prion disease model and was active in the *in vivo* tunicamycin assay. In this study of chronic treatment with GSK2606414 in SOD1G93A mice, several statistical analysis methods showed that both male and female mice treated with GSK2606414 experienced a marked decrease in body weight maintenance (S2 Fig). The progression of neurological disease tended to be faster by about 3 days in female mice treated with GSK2606414, as indicated by ordinal logistic regression analysis, though the effect was not statistically significant (p=0.38). In contrast, male mice tended to experience a slower progression, by about three days on average, though also non-significantly (p=0.11). When the genders were combined there was no net effect on neurological score progression (Fig 5A). Disease onset was variably shortened in female, but variably prolonged in male GSK2606414-treated animals. The survival duration was not impacted in female mice, but there was a tendency towards a non-significant increase in male mice. When the genders were combined there was no net beneficial effect on the time to onset of definitive neurological disease, or on survival duration (Fig 5B, Table 1). The high variability among the drug-treated mice of both genders, in combination with the observed significant weight loss, suggests that the treatment with GSK2606414 was not well-tolerated by some of the mice.

**Fig 5A.**
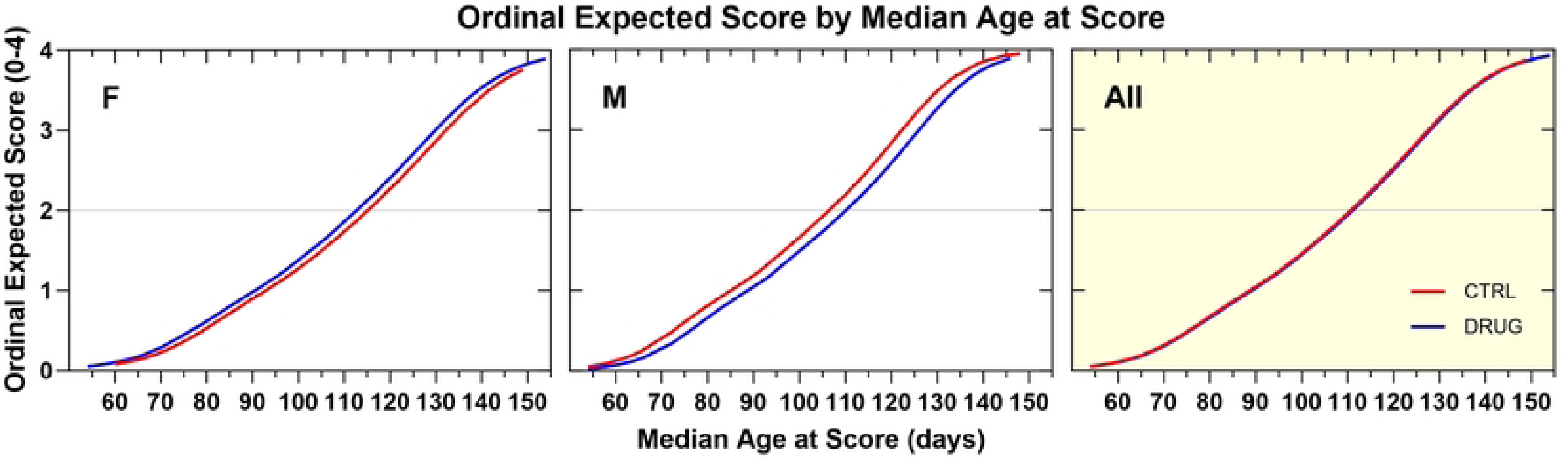
Ordinal logistic regression analyses of the effect of GSK2606414 50 mg/kg, PO, QD on Neurological Disease Progression. Female GSK2606414-treated animals tended to show a faster rate of progression that vehicle controls by about 3 days (p=0.38). Male treated mice tended to progress more slowly than vehicle controls by about 3 days (p=0.11). Neither effect was statistically significant. There was no effect on progression when the genders were combined.

**Fig 5B.**
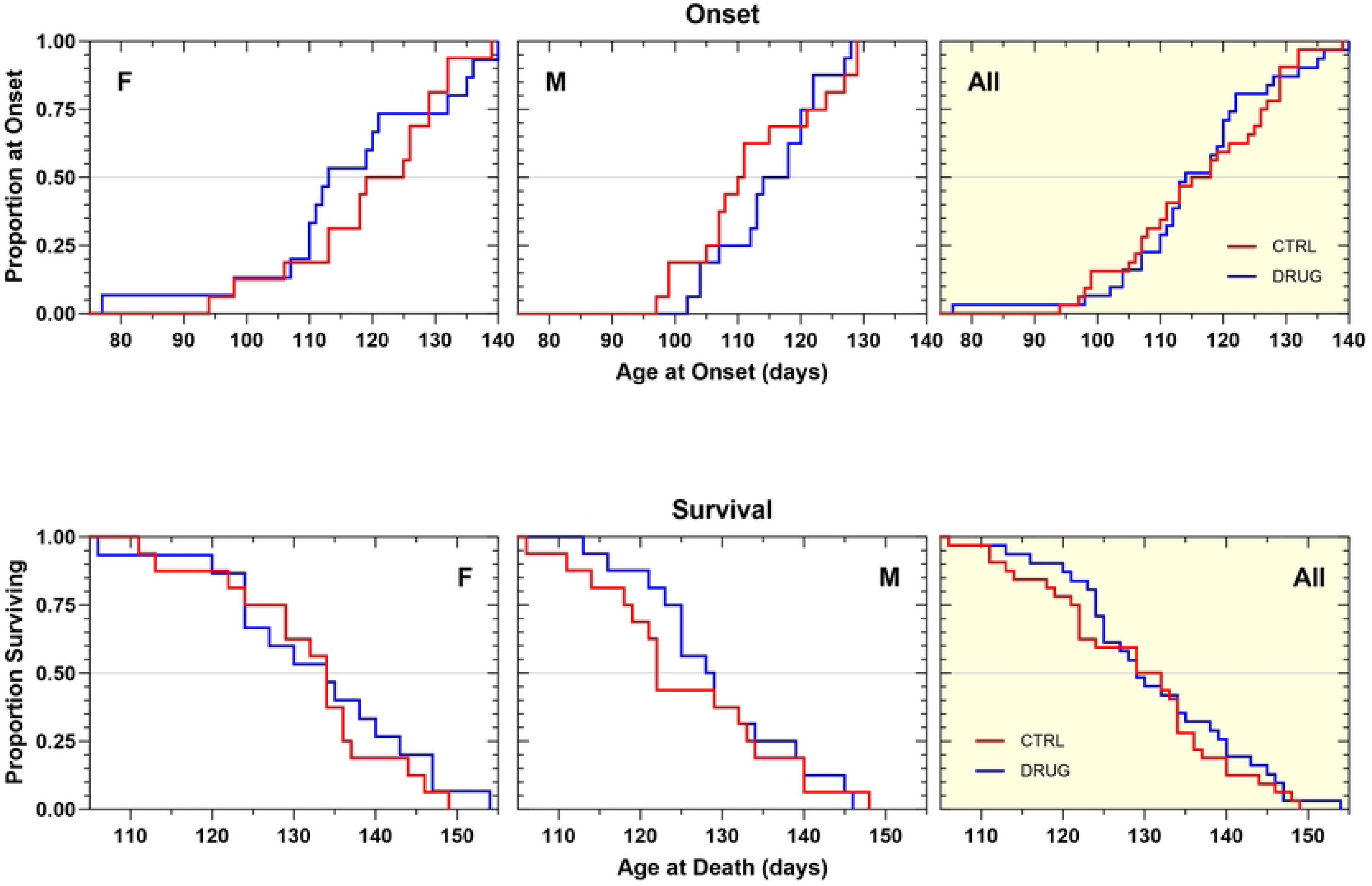
Kaplan–Meier analyses of the effect of GSK2606414 50 mg/kg, PO, QD on Disease Onset (top panels) and Survival Duration (bottom panels). Time to onset of definitive neurological disease was variably shortened in female GSK2606414-treated animals compared to vehicle control animals (-9 d, NS). Male GSK2606414-treated animals tended to have a variable delay in onset (+5.5 d, NS). When the genders were combined age at onset of GSK2606414-treated animals was similar to controls. Survival duration in GSK2606414-treated female animals was identical to vehicle control animals.

**Table 1.** Summary of survival efficacy study outcomes.

|  | **GSK2606414** |  |  | **Salubrinal** |  | **Sephin1** |  | **Clonidine** |
| --- | --- | --- | --- | --- | --- | --- | --- | --- |
| **Regimen** | 50mg/kg, PO, qd | 50mg/kg in chow, ad lib | 18 mg/kg, PO, qd | 5mg/kg, IP, qd | 15mg/kg, IP, qd | 4mg/kg, IP, qod | 10mg/kg, IP, qd | 0.5mg/kg, IP, qod |
| **Body Weight** | Worsened (p=0.001) | Worsened (p=0.0001) | No change | Improved (p=0.03) | No change | No change | No change | No change |
| **Progression** | No change<sup>a</sup> | No change (accelerated in males by 7 days p=0.009) | No change | No change | No change | No change | No change | No change (decelerated in females by 4 days p=0.03) |
| **Onset** | No change<sup>a</sup> | No change<sup>b</sup> | No change | No change | No change | No change | Delayed by 5 days (p=0.04) | No change |
| **Survival** | No change<sup>a</sup> | No change<sup>b</sup> | No change | No change | No change | No change | No change | No change (extended in females by 8 days p=0.05) |

Male GSK2606414-treated tended to show extended survival (+6.5 d, NS). There was no net effect on survival duration.

In a subsequent survival efficacy study, GSK2606414 was formulated in standard chow and was made available to mice *ad libitum*, to provide 50 mg/kg/day, based on a 4 g average daily chow consumption (previously determined). This method of administration, associated with food consumption, was selected to produce low peak concentrations in blood that could avoid tolerability problems during chronic treatment and yet have the drug consistently available in circulation. The mice, on average, consumed equal amounts of the GSK2606414-treated chow compared to standard chow indicating that there was no taste aversion associated with this formulation. Like the results of the previous survival efficacy study, treatment with GSK2606414 negatively impacted body weight maintenance in this chronic study. GSK2606414-treated animals showed a body weight decrease of 1.2-1.5 g (p<0.0001) when compared to vehicle-treated animals (S3 Fig). Unlike the previous study, GSK2606414 treatment significantly accelerated the rate of neurological disease progression (Fig 6A) and shortened survival duration in male mice (Fig 6B) and not the female. GSK2606414 treatment did not produce significant changes in female mice or when the two genders were analyzed together (Fig 6; Table 1). Overall, this treatment regimen of GSK2606414 did not produce a clear therapeutic benefit in the SOD1G93A mouse model of ALS. On the contrary, there was evidence of an increase in disease severity in male mice.

**Fig 6A.**
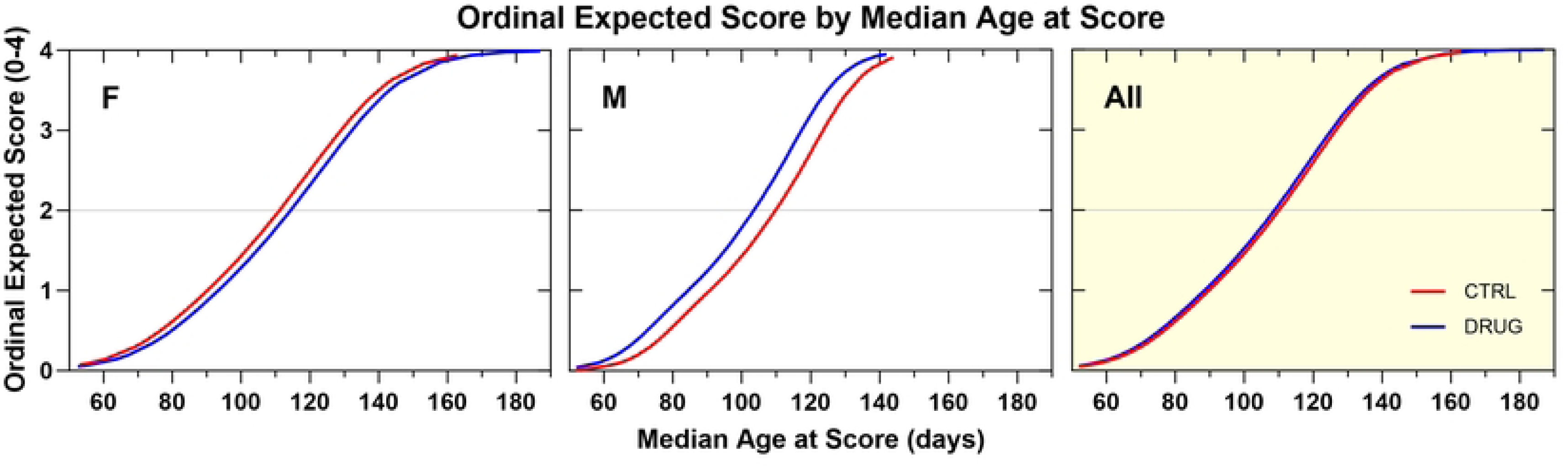
Ordinal logistic regression analyses of the effect of GSK2606414 50 mg/kg in Standard Chow, ad libitum on Neurological Disease Progression. GSK2606414-treated females tended to progress more slowly by about 4 days compared to vehicle control females (NS). Male GSK2606414-treated animals progressed more quickly than controls by about 6 days (p=0.009).

**Fig 6B.**
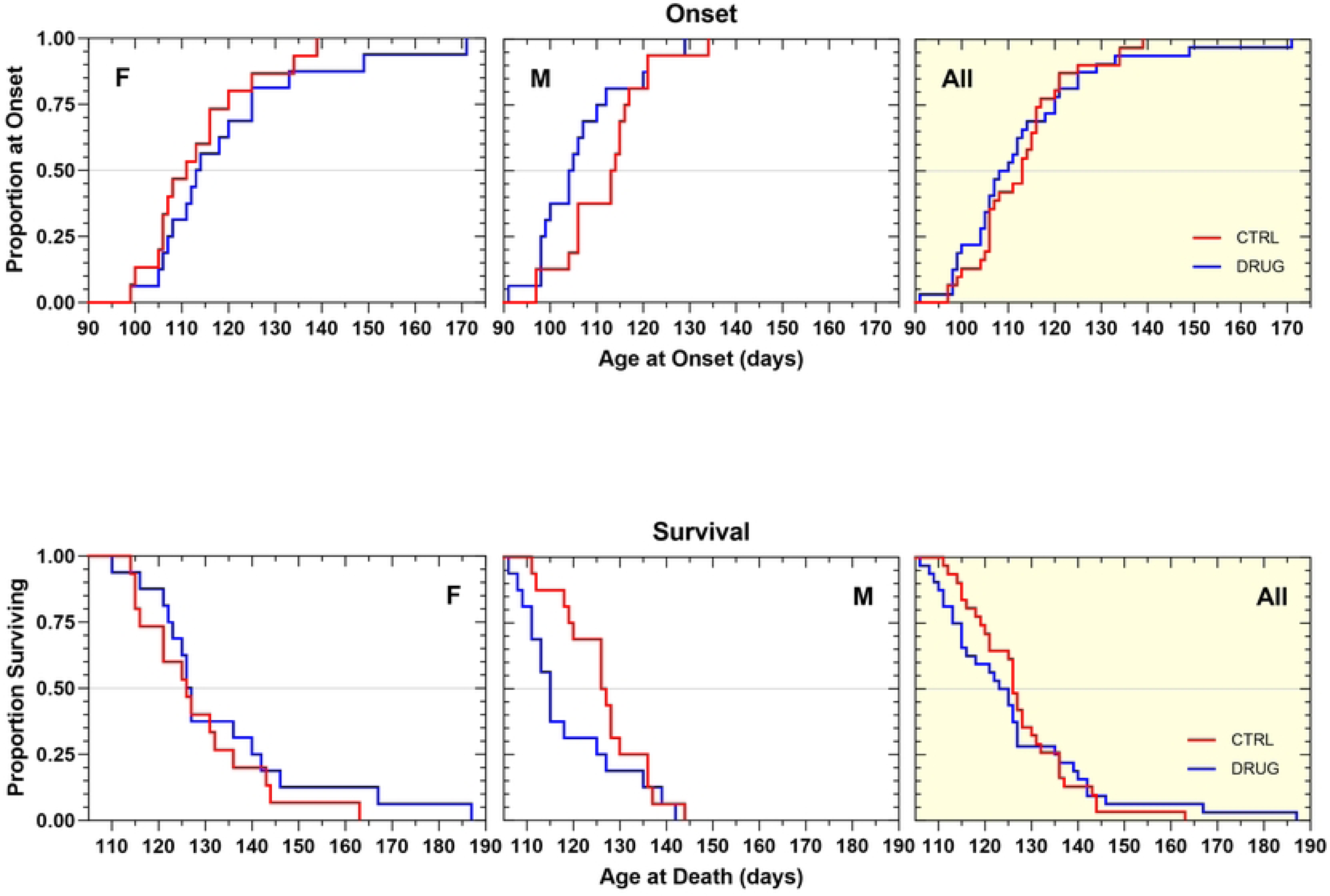
Kaplan–Meier analyses of the effect of GSK2606414 50 mg/kg/d in Standard Chow, ad libitum on Disease Onset (top panels) and Survival Duration (bottom panels). Time to disease onset tended to be later in female (+2.5 d, p=0.11) and earlier in male (-9 d, p=0.15) GSK2606414-treated animals. When the genders were analyzed together there was no net effect. Survival duration tended to be shorter (-12 d, p=0.12) in male, but not in female GSK2606414-treated animals. There was no overall change in disease onset or survival duration as a result of treatment.

In our final attempt to elicit a beneficial effect of the drug, we tested an approximately 3-fold lower dose of GSK2606414 (18 mg/kg) via daily oral gavage, the rationale being that any potential beneficial effects of the drug in the previous studies were compromised by drug toxicity. The 18 mg/kg dose had also produced a significant effect in the *in vivo* tunicamycin assay. This treatment regimen did not produce a clear beneficial effect on body weight maintenance (S4 Fig), did not delay the rate of neurological disease progression (Fig 7A), had no effect on the onset of definitive disease and did not extend survival in either gender (Fig 7B; Table 1). Overall, there was no clear evidence of therapeutic benefit with this treatment regimen, but also no clear evidence of any increase in disease severity.

**Fig 7A.**
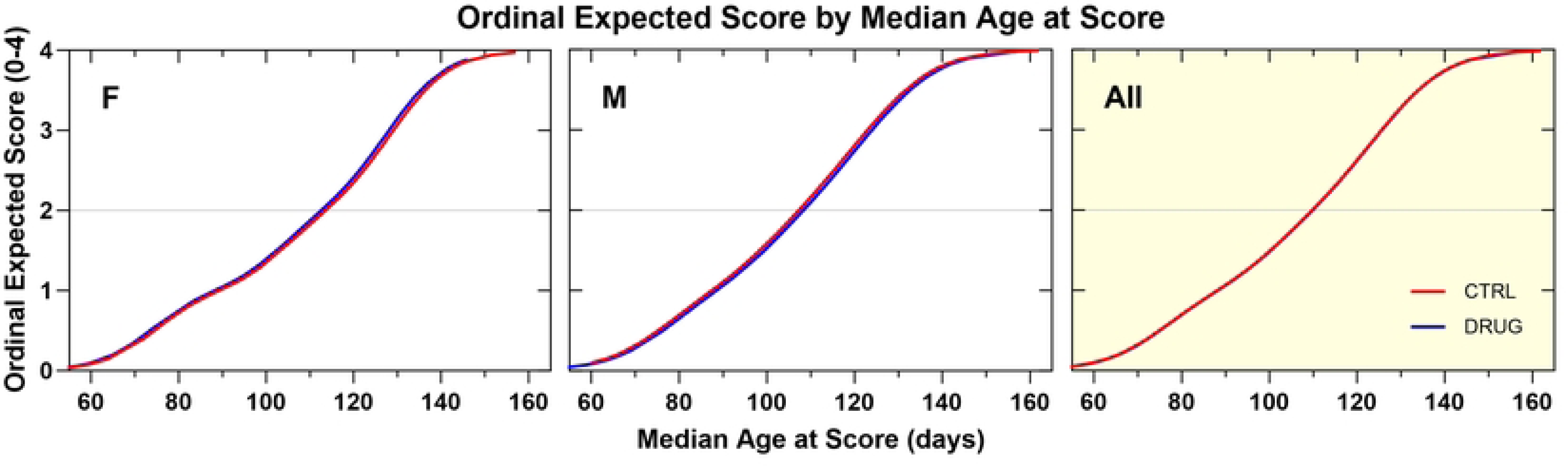
Ordinal logistic regression analyses of the effect of GSK2606414 18 mg/kg, PO, QD on Neurological Disease Progression. GSK2606414-treated animals, in both genders, progressed in a manner similar to controls.

**Fig 7B.**
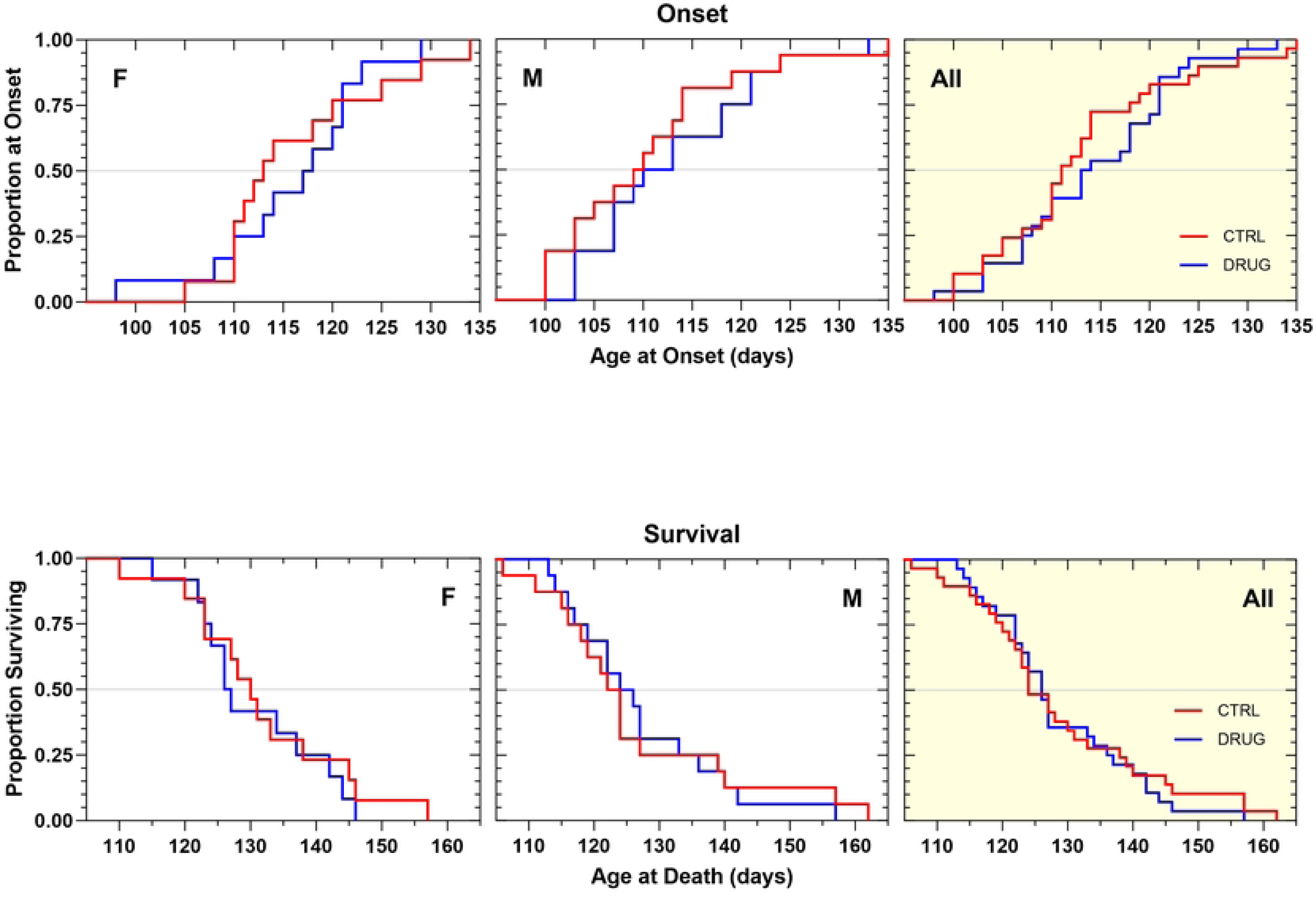
Kaplan–Meier analyses of the effect of GSK2606414 18 mg/kg, PO, QD on Disease Onset (top panels) and Survival Duration (bottom panels). Kaplan–Meier analysis of the effect of GSK2606414 on disease onset (top panels) and survival duration (bottom panels). Data are represented as females only, males only, and combined (ALL). There was no overall change in disease onset or survival duration as a result of treatment.

Notably, none of the three GSK2606414 treatment regimens that we investigated had an impact on terminal blood glucose levels as previously reported by others (data not shown).

### Salubrinal Survival Efficacy Studies

Despite not exhibiting any activity in the *in vivo* tunicamycin assay, we tested salubrinal in efficacy studies, due to previous reports of positive findings in SOD1G93A mice^24^. The positive effects of salubrinal were reported by only one group and have not been validated by others. This could be due to salubrinal’s poor pharmacological properties, particularly its low solubility, which posed challenges in administering it safely to animals over a prolonged period. Thus, it was important for us to develop formulations for intraperitoneal salubrinal administration that were well-tolerated for chronic administration. After testing several excipients in preliminary preclinical formulation studies, we arrived at a formulation consisting of 5% N-Methyl-2-pyrrolidone (NMP) and 5% Kolliphor® HS15 in filtered tap water that was deemed to be safe in two-week long tolerability studies.

Salubrinal was tested at two concentrations, 5 and 15 mg/kg, in two different standardized litter-matched and gender-balanced drug efficacy studies in SOD1G93A mice (Figs 8,9). The 5 mg/kg dose was chosen to approximate the same drug exposure as in the published study that had shown benefits, and in the subsequent study we administered a three-fold higher dose to see whether increasing the dose improved outcomes. Apart from a small, but significant (p=0.03), improvement in the median body weight change (S5, S6 Figs), which was only present in the 5 mg/kg cohort, neither chronic salubrinal dosing regimen beneficially affected the rate of neurological disease progression, disease onset, or survival duration (Figs 8,9; Table 1). It is important to note that no adverse effects of salubrinal were observed.

**Fig 8A.**
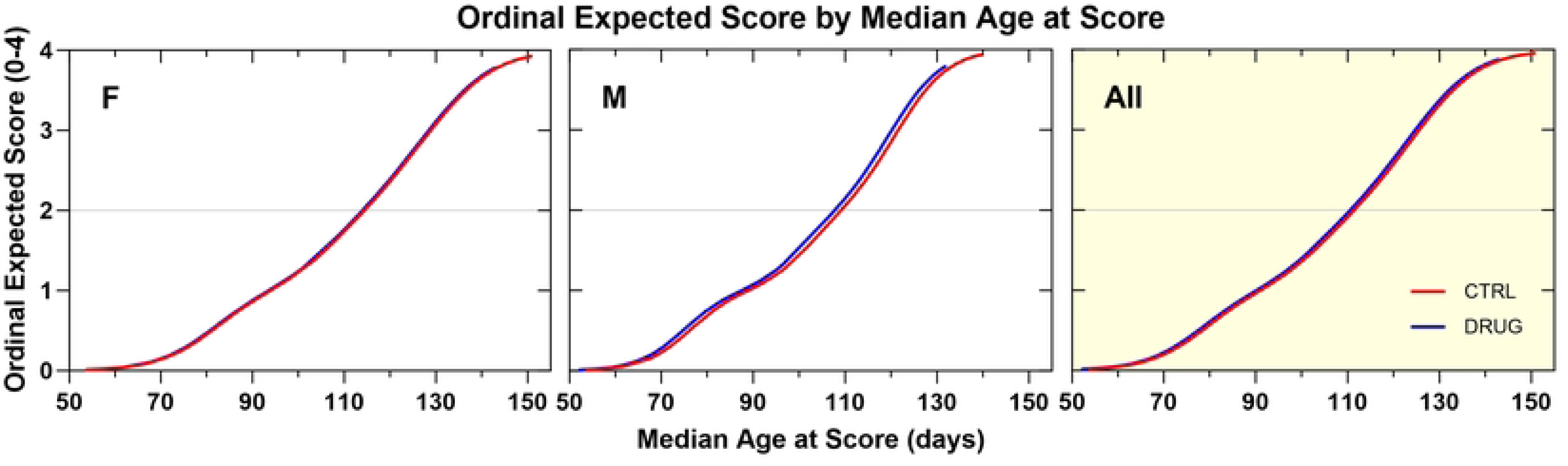
Ordinal logistic regression analyses of the effect of salubrinal 5mg/kg, IP, QD on Neurological Disease Progression. A similar rate of progression was observed in salubrinal-treated animals compared to vehicle control animals.

**Fig 8B.**
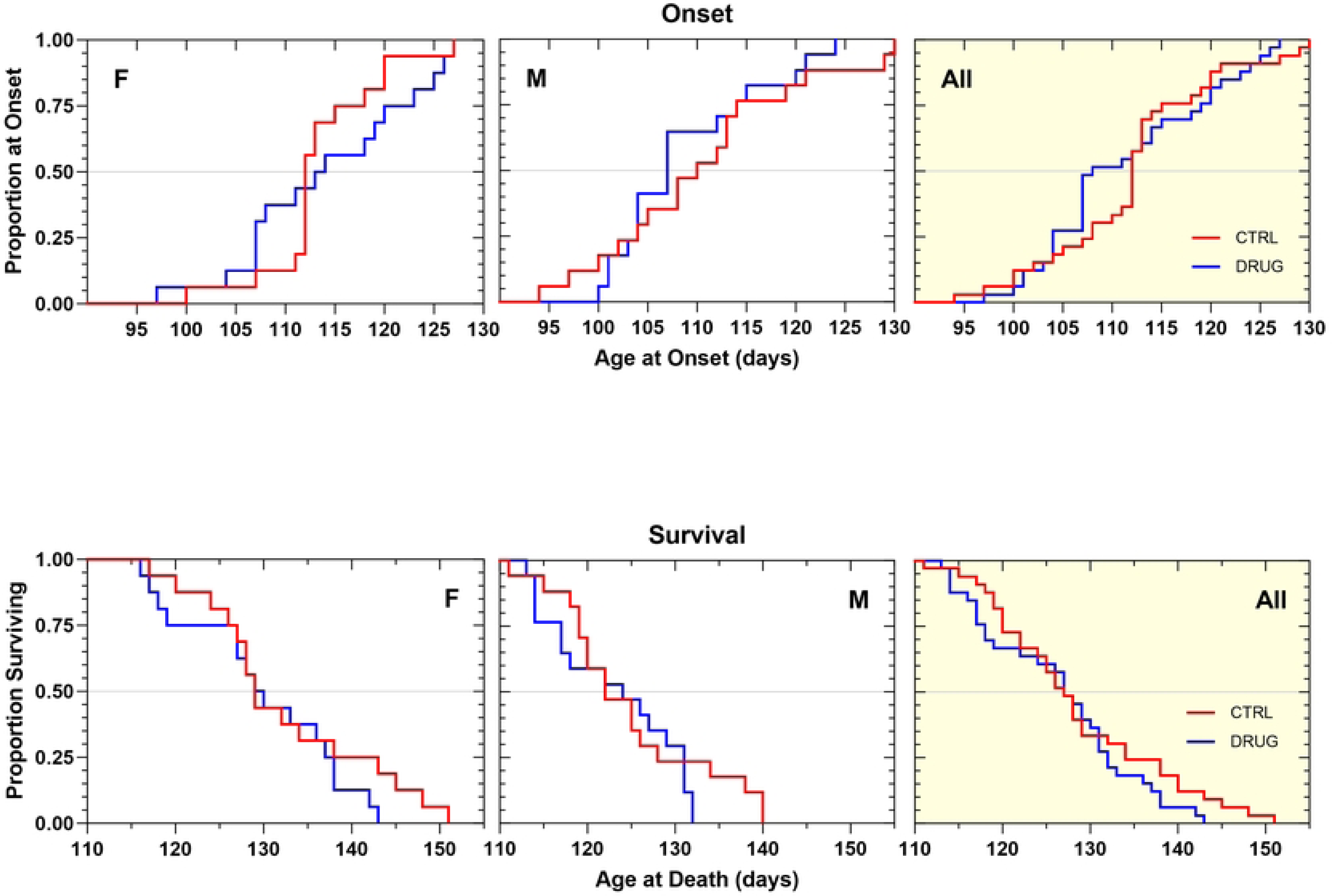
Kaplan–Meier analyses of the effect of salubrinal 5 mg/kg, IP, QD on Disease Onset (top panels) and Survival Duration (bottom panels). Time to disease onset was similar in salubrinal-treated and vehicle control animals. Survival duration was similar in salubrinal-treated and vehicle control animals.

**Fig 9A.**
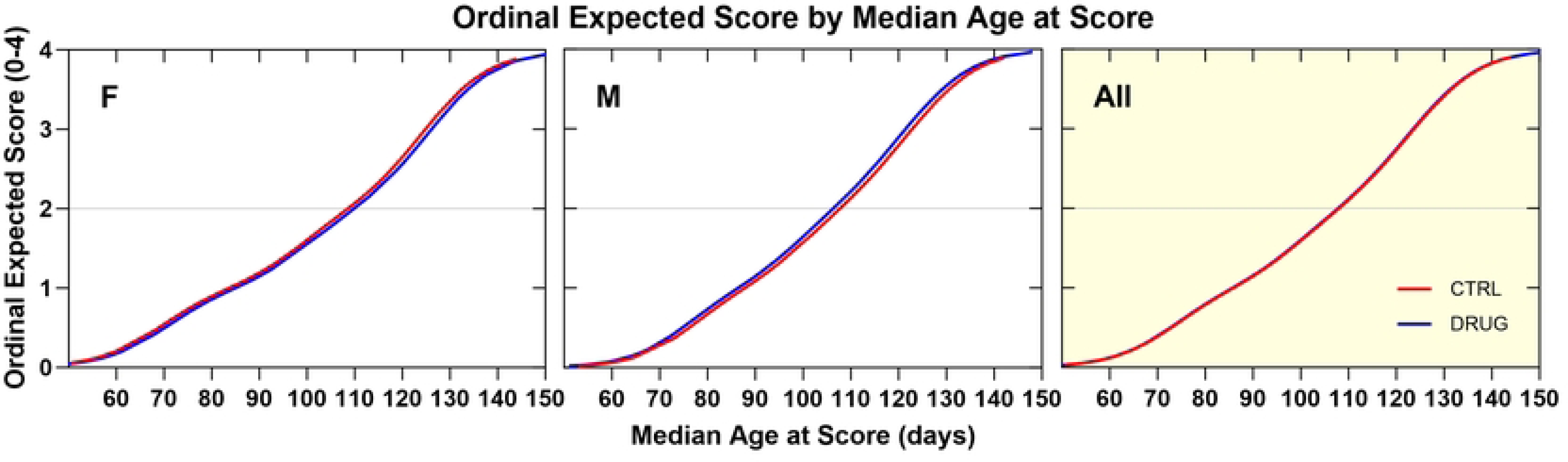
Ordinal logistic regression analyses of the effect of salubrinal 15mg/kg, IP, QD on Neurological Disease Progression. A similar rate of progression was observed in salubrinal-treated animals compared to vehicle control animals.

**Fig 9B.**
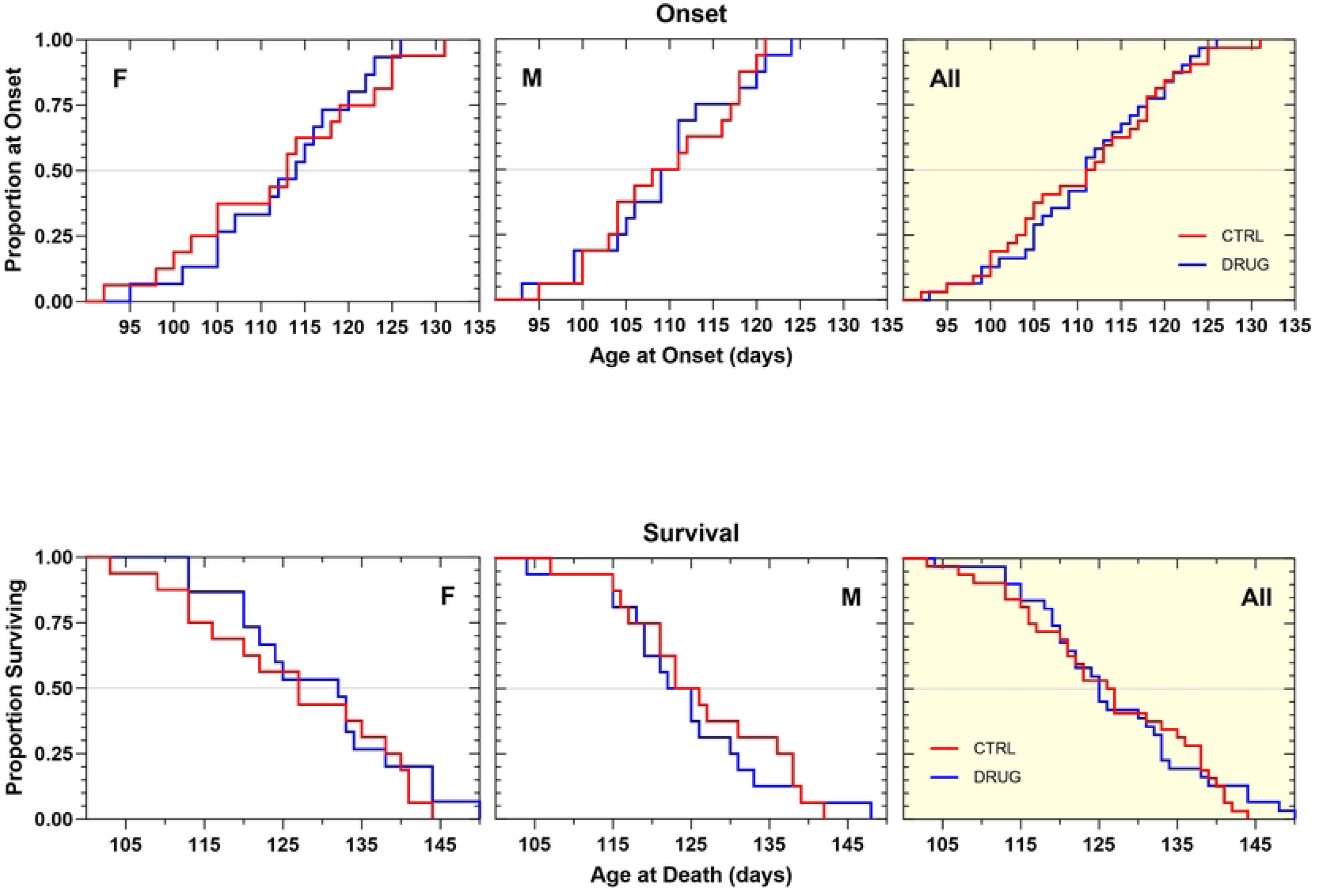
Kaplan–Meier analyses of the effect of salubrinal 15 mg/kg, IP, QD on Disease Onset (top panels) and Survival Duration (bottom panels). Time to disease onset was similar in salubrinal-treated and vehicle control animals. Survival duration was similar in salubrinal-treated and vehicle control animals.

### Sephin1 Survival Efficacy Studies

We then proceeded to test Sephin1, a very close analog of the previously tested guanabenz, that lacks alpha-2-adrenergic agonist effects. This study sought to determine whether an otherwise identical compound and dosing regimen could produce beneficial effects on ALS-related proteostasis when alpha 2 adrenergic receptor agonism-related side effects were eliminated. Sephin1 was tested twice, in standardized litter-matched and gender-balanced drug efficacy studies in SOD1G93A mice. In the first study, it was tested at 4 mg/kg, dosed every other day, intraperitoneally to match the dose of guanabenz previously tested in the same animal model^50^. In our hands, this dosing regimen did not produce an overall therapeutic benefit, though the adverse side-effects associated with guanabenz treatment were not observed. Sephin1-treated animals’ body weights were slightly and variably improved by treatment (S7 Fig). Disease onset, rate of neurological disease progression, and survival duration were not beneficially affected by chronic treatment with this dosing regimen (Figs 10A, 10B; Table 1).

**Fig 10A.**
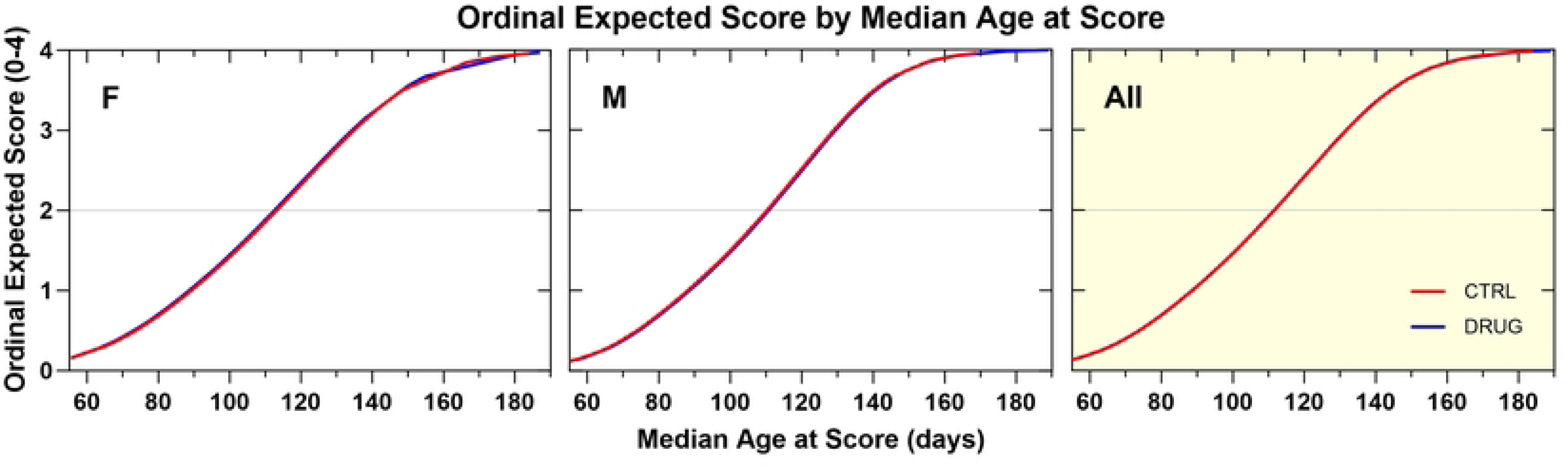
Ordinal logistic regression analyses of the effect of Sephin1 4 mg/kg, IP, QOD on Neurological Disease Progression. Sephin1-treated animals progressed at the same rate as vehicle-treated animals.

**Fig 10B.**
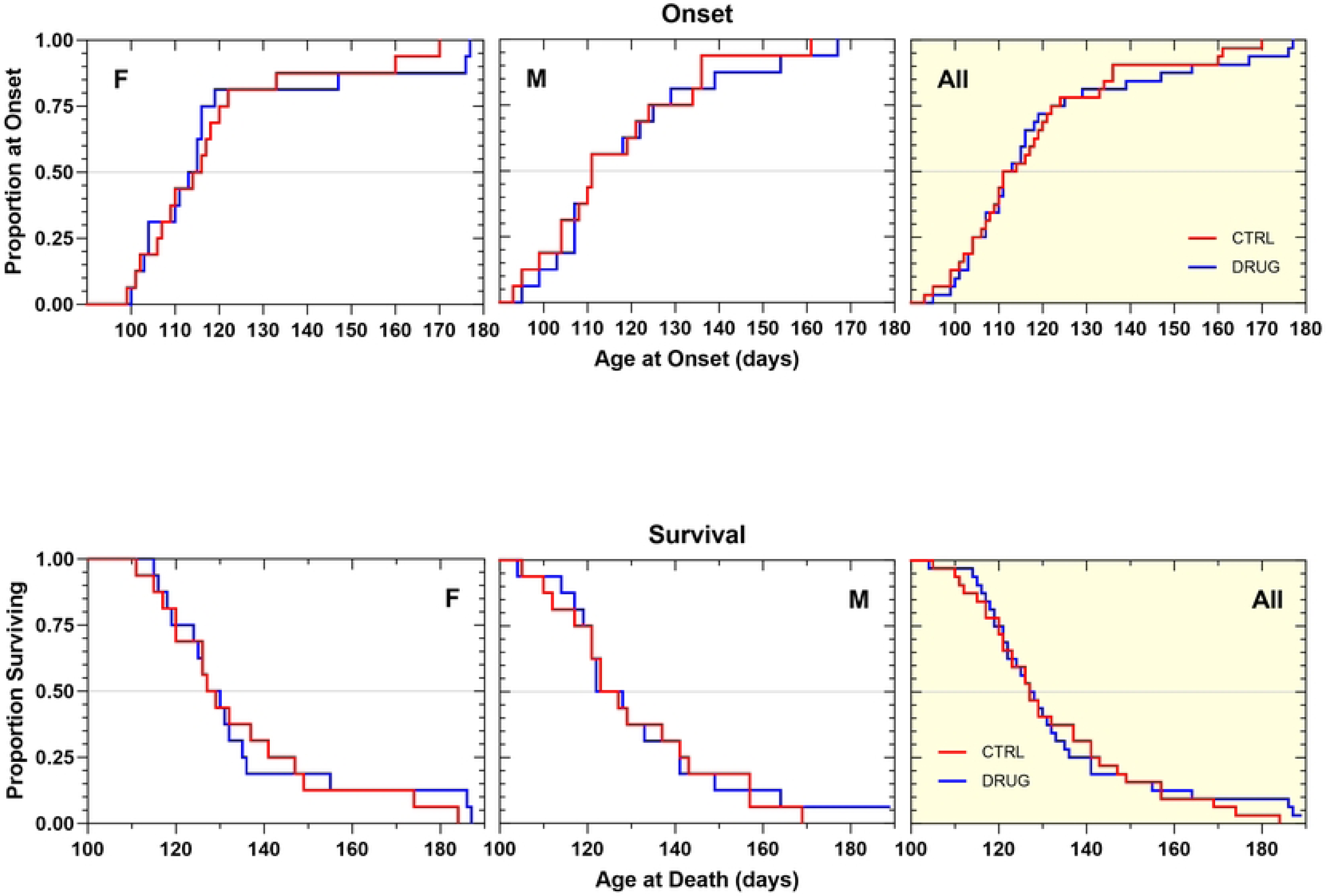
Kaplan–Meier analyses of the effect of Sephin1 4 mg/kg, IP QOD on Disease Onset (top panels) and Survival Duration (bottom panels). There was no overall change in disease onset or survival duration as a result of treatment.

The second efficacy study with Sephin1 began with a dose of 15 mg/kg administered intraperitoneally, a dose that was determined to be efficacious in the *in vivo* tunicamycin assay. However, after three days of administration at this dose, the mice experienced rapid body weight loss. As a result, the dose was reduced to 10 mg/kg, effectively stopping the rapid weight loss and causing no additional drug-related adverse effects for the rest of the study period. Neurological score progression tended to be slightly slower in Sephin1-treated animals (Fig 11A) (+3 d, *p=0.10*). This treatment regimen produced a statistically significant therapeutic benefit on disease onset (Fig11B; Table 1), which was delayed in Sephin1-treated animals compared to vehicle controls by 5 days (p=0.05). Additionally, survival duration was extended by a similar amount, but this effect was not statistically significant. Body weight tended to be less well maintained by several measures in Sephin1-treated animals, although most of the effect was contributed by males (S8 Fig).

**Fig 11A.**
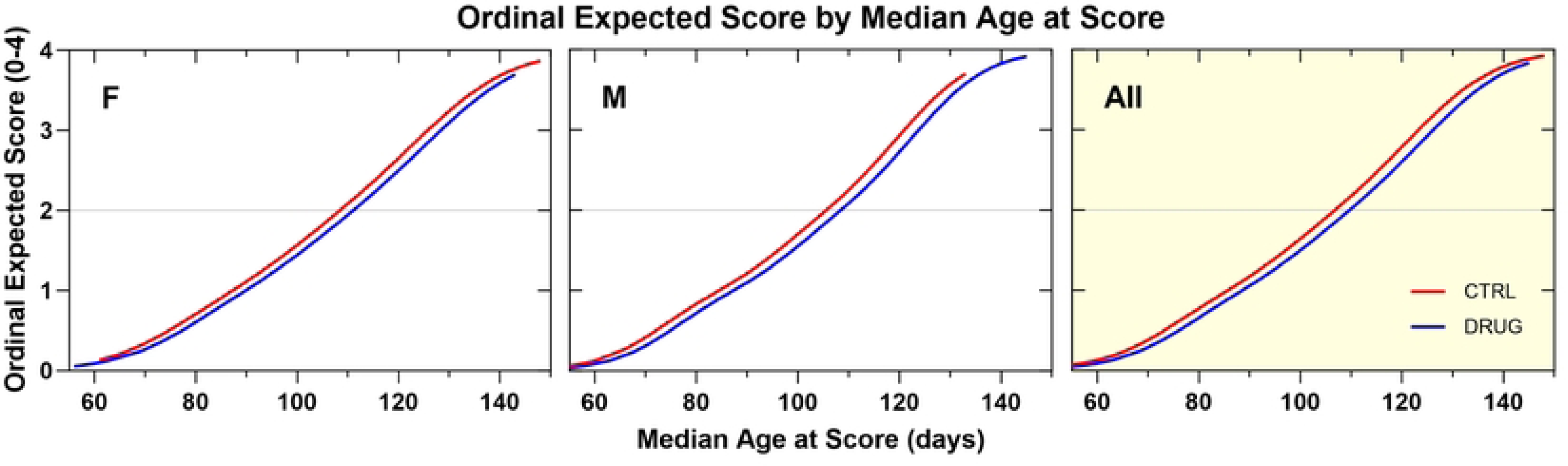
Ordinal logistic regression analyses of the effect of Sephin1 10 mg/kg, IP, QD on Neurological Disease Progression. Sephin1-treated animals of both genders showed a slightly slower rate of progression compared to vehicle control animals by about 3 days (NS).

**Fig 11B.**
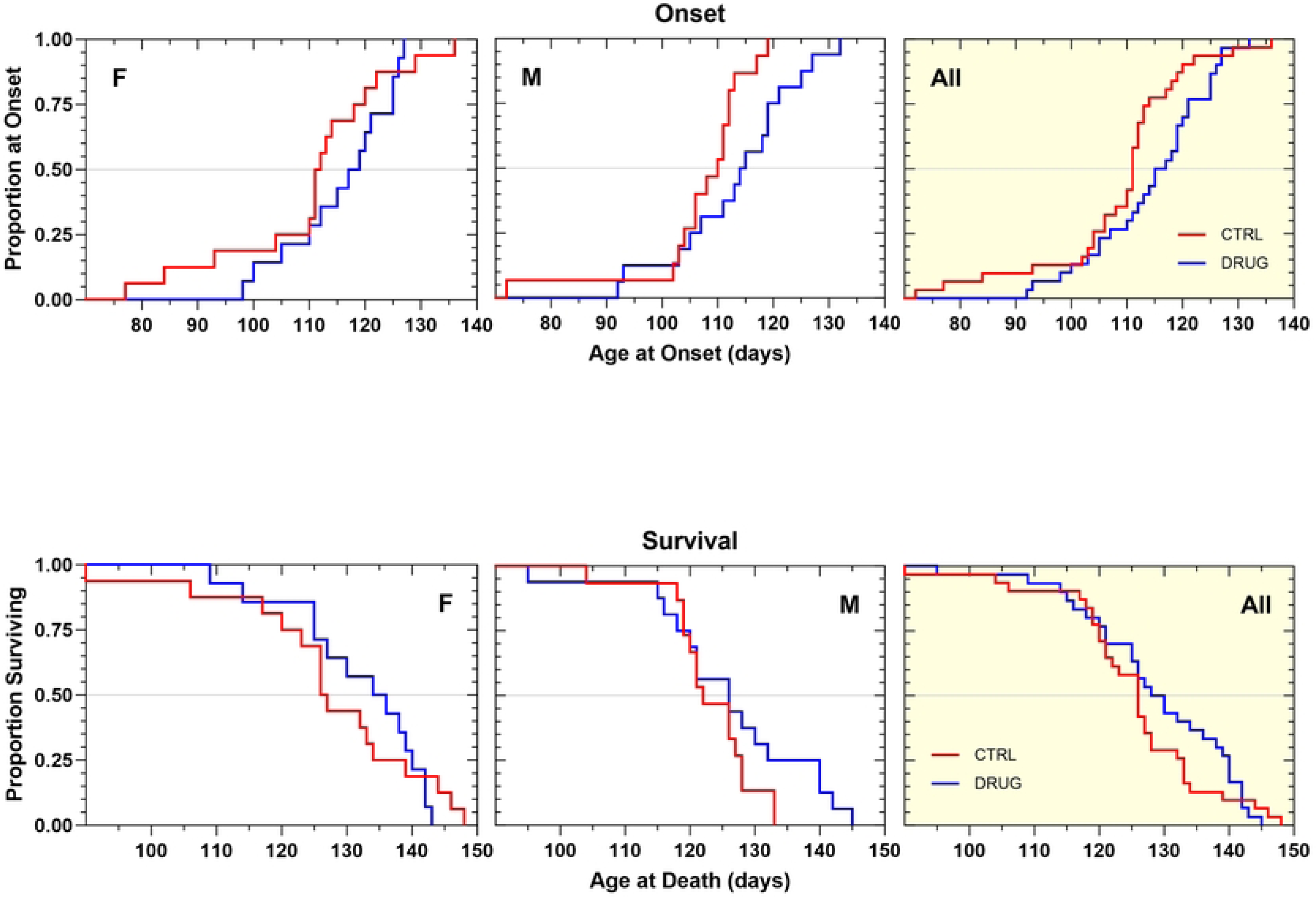
Kaplan–Meier analyses of the effect of Sephin1 10 mg/kg, IP, QD on Disease Onset (top panels) and Survival Duration (bottom panels). There was a statistically significant overall delay in disease onset by 5 days (p=0.05) as a result of treatment. Survival duration tended to be extended by 3 days, though the effect was variable and not statistically significant.

### Clonidine Survival Efficacy Study

To directly test whether alpha-2-adrenergic receptor agonism exacerbated the disease progression in SOD1G93A mice, we treated mice with the potent alpha-2-adrenergic agonist class prototype drug clonidine, given as the water-soluble hydrochloride salt dissolved in normal saline. Clonidine, which distributes very well to the central nervous system, was tested at 0.5 mg/kg, dosed every other day, intraperitoneally in a standardized litter-matched and gender-balanced drug efficacy study. This dose was chosen to approximate the alpha-2-adrenergic receptor agonism that would have been caused by guanabenz in our previous studies, based on reports comparing the efficacy and potency of several alpha-2-adrenergic receptor agonists and antagonists in radioligand binding^51^ and phenotypic assays^52^.

To our surprise, clonidine not only did not worsen the condition of SOD1G93A mice, but in female mice there were also signs of a therapeutic benefit (Table 1). In female clonidine-treated animals there was an improvement in body weight maintenance (S9 Fig), a significant slowing of disease progression (Fig 12A, +4 d, p=0.03) and prolongation of survival duration (Fig 12B, + 8 days, p=0.05) compared to controls. Male mice trended towards the opposite effects, so that when two genders were combined any beneficial effects on survival duration, the time to onset of definitive neurological disease, and the neurological score progression were neutralized.

**Fig 12A.**
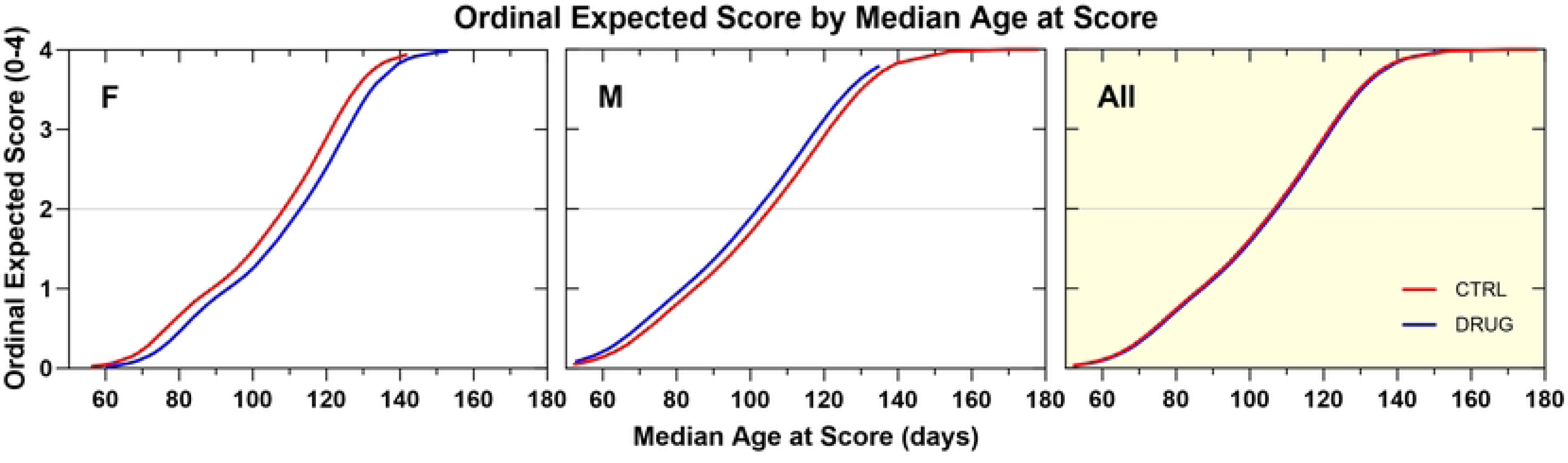
Ordinal logistic regression analyses of the effect of Clonidine 0.5 mg/kg, IP, QOD on Neurological Disease Progression. Clonidine-treated females showed a statistically significantly slower progression (+4 d, p=0.03), and males tended to show a more rapid progression (-4 d, p=0.17). Treated animals progressed in a manner essentially identical to controls when both genders were combined.

**Fig 12B.**
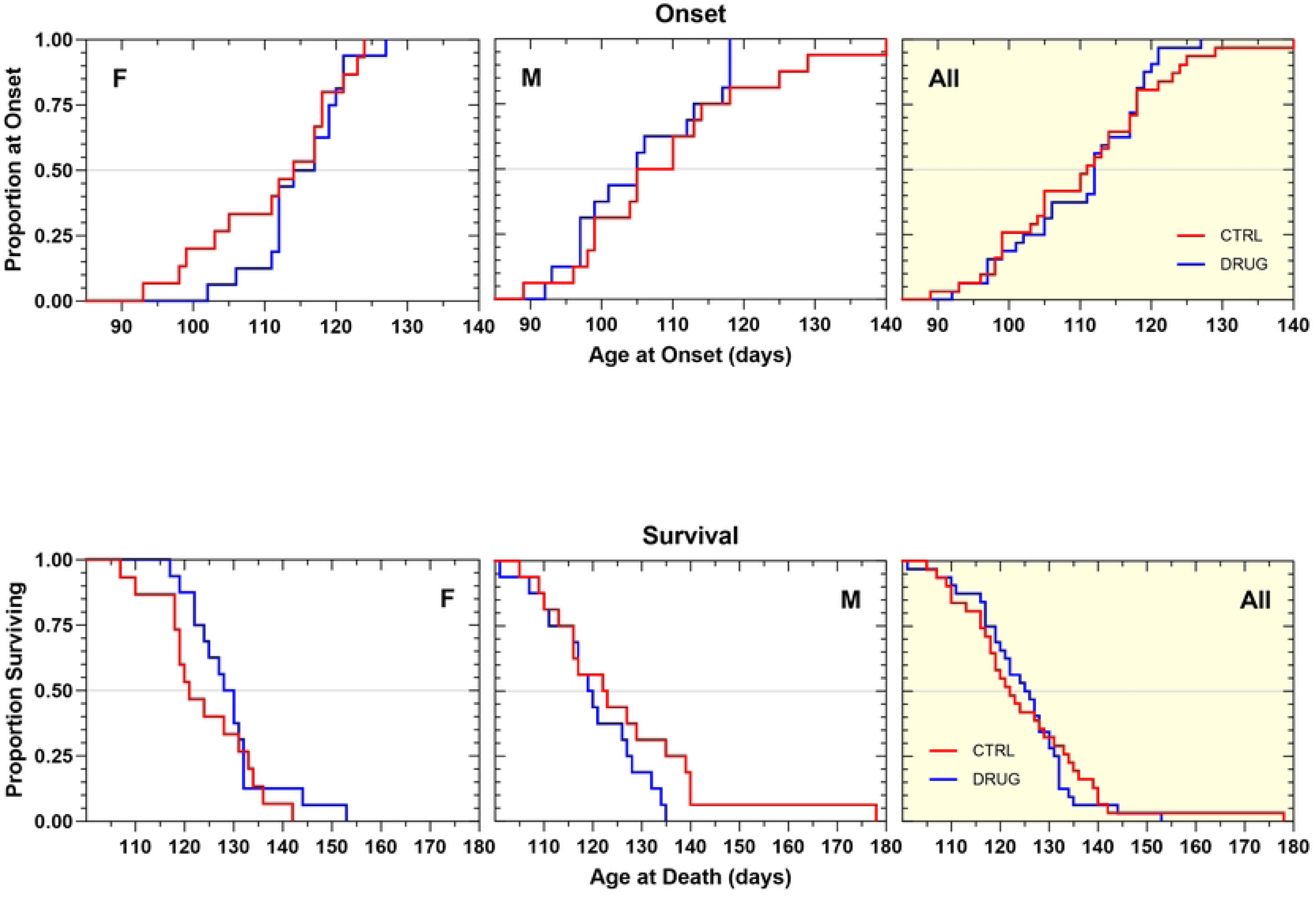
Kaplan–Meier analyses of the effect of Clonidine 0.5 mg/kg, IP, QOD on Disease Onset (top panels) and Survival Duration (bottom panels). There was no overall change in disease onset or survival duration as a result of treatment. Survival duration was significantly prolonged in clonidine-treated female animals compared to female vehicle control animals by 8 days (p=0.05).

## Discussion

For more than two decades, the PERK branch of the UPR has been considered a potential drug target for ALS. However, attempts to develop therapies through pharmacological modulation of PERK have so far been unsuccessful. While some preclinical studies have shown promise, overall, preclinical outcomes have been inconsistent and conflicting. The reasons for this remain unclear. Could it be that dosing regimens for the drugs tested were suboptimal? Were the preclinical study designs flawed? Or is the PERK pathway so complex that developing small molecule modulators is extremely challenging? In our studies we sought to determine the utility of PERK as an ALS drug target. To reconcile the contradictory results of previous research, we evaluated three commonly used PERK modulators - GSK2606414, salubrinal, and Sephin1 - in the same ALS-relevant screening assay and animal model. To assess their ability to modulate the PERK pathway and gain insight into their mechanisms of action, we developed a mouse-based assay that modeled UPR pathway activation using tunicamycin. To evaluate their therapeutic potential, we tested them for efficacy in the SOD1G93A mouse model of ALS.

The results of the mouse-based tunicamycin assay largely confirmed the PERK-modulatory properties of these compounds, though there were some novel findings that may have implications for future drug development efforts. Specifically, we found that the specific PERK kinase inhibitor GSK2606414 attenuated the expression of PERK pathway genes such as ATF4 and GADD34, which serve to disinhibit protein translation, and at the same time decrease CHOP activation avoiding pro-apoptotic pathways. GSK2606414 also attenuated the expression of ATF6 and its downstream targets, the ER chaperones HSPA5, EDEM1, DNAJb9, via non-canonical signaling that has not previously been described. As anticipated, based on its mechanism of action, GSK2606414 did not have a modulatory effect on the IRE1α pathway and downstream targets. On the other hand, salubrinal, a PERK activator, did not influence any of the activated UPR genes we looked at. This was consistent with its mechanism of action since it indirectly enhances eIF2α phosphorylation, by inhibiting GADD34-mediated dephosphorylation of eIF2α, leading to prolonged activation of the UPR response. Since in this assay, the UPR response was already maximally activated, a salubrinal-induced activation was largely masked. For multiple transcripts assayed, salubrinal treatment tended to elicit expression levels greater than those induced by tunicamycin alone, though these effects were variable and not statistically significant. Finally, Sephin1, another PERK pathway activator we tested, produced some unexpected outcomes. It decreased the upregulation of CHOP, EDEM1, DNAJb9, and HSPA5, but did not affect the expression of the ATF4 gene, possibly indicating that its modulatory effects were not mediated via an inhibition of the PERK pathway. Instead, it is likely that it was Sephin1’s actions on the IRE1α pathway that mediated these effects, as suggested by a trend towards an attenuation in the expression of Ire1α (p=0.0946).

The mouse-based tunicamycin assay used herein provided insights into the mechanism of action of the UPR modulators, though it also had several limitations. The signature of genes upregulated by TUN was similar to that of ALS, in both patients and animal models, but the extent of upregulation produced by TUN was excessive and surpassed even pathological levels. In hindsight, different assay parameters could have been chosen, by using lower concentrations of TUN to produce a more modest upregulation of UPR genes. This may have enabled us to detect drug-induced enhancements rather than just attenuations. Additionally, the use of liver as a surrogate tissue for spinal cord and brain requires that the relevant pharmacological and pharmacokinetic properties of drugs destined for CNS disorders need to be determined beforehand. Finally, this assay, given its laborious nature and the requirement for many mice, is not suited as a primary, high-throughput screen, but nevertheless can be employed as a valuable secondary target engagement screen for UPR-active compounds.

GSK2606414 unequivocally failed to result in a therapeutic benefit in three different survival efficacy studies in SOD1G93A mice despite its ability to inhibit PERK and its downstream targets in the *in vivo* tunicamycin assay. There were even signs to suggest that it worsened the clinical condition of SOD1G93A mice by causing significant weight loss and variably decreasing the survival duration. Moreover, the blood glucose levels in SOD1G93A mice, treated with GSK2606414 over a prolonged period, were normal, suggesting that on-target pancreatic toxicity effects of the drug were unlikely. We interpret these results to mean that inhibition of the PERK pathway reduces the ability of mice to respond to the increased amount of misfolded SOD1 to restore homeostasis. Our findings support the results of several genetic studies where the PERK pathway was manipulated and the disease was accelerated, however they contradict the outcomes of a pharmacological study that reported GSK2606414-induced neuroprotection in a prion disease mouse model^27^. In the prion disease model however, ER stress was induced by a single inoculation of prion containing-brain homogenates and the level of PERK activation was likely well below that of the transgenic models where there is a continuous production of misfolded proteins.

Salubrinal, an enhancer of the PERK pathway, did not have any impact on disease onset, rate of disease progression, or survival duration in two different survival efficacy studies conducted in the SOD1G93A model, which was consistent with its lack of activity in the *in vivo* tunicamycin assay. This result suggests that the PERK pathway was already optimally activated in SOD1G93A mice as a compensatory mechanism to counteract protein dyshomeostasis that was already in motion, and any additional effect of salubrinal was insufficient to produce a therapeutic benefit. Since salubrinal did not cause any adverse reactions, and in one of the studies produced a small improvement in body weight maintenance, one can speculate that higher doses of salubrinal could have produced a therapeutic benefit, though because of its insolubility we were unable to test that hypothesis. It is worth noting that our findings differ from a previous study in SOD1G93A mice, which claimed that salubrinal extended survival^25^. However, we consider the results of that study to be unreliable, as there were only four mice in each experimental group.

The next molecule we tested in the SOD1G93A model was Sephin1, which is believed to enhance the activity of the PERK pathway by prolonging the phosphorylation of eIF2α, like salubrinal does. However, Sephin1 works by inhibiting GADD34, a regulatory subunit of protein phosphatase 1 that dephosphorylates eIF2α, rather than directly suppressing the phosphatase complexes that dephosphorylate eIF2α. We conducted two survival efficacy studies with Sephin1. In the first study, we used a treatment regimen that we had previously used for its close analog guanabenz, in which Sephin1, like guanabenz, did not provide any therapeutic benefits. In the second survival efficacy study, we increased the dose level of Sephin1 that we tested to 10mg/kg and with daily dosing, which resulted in notable improvements in the clinical condition of the SOD1G93A mice, with some therapeutic indicators being statistically significant. Our findings validate the outcomes of several other preclinical studies that demonstrated Sephin1’s neuroprotective properties^35,36,37^. However, since Sephin1 did not affect the expression of ATF4, our findings challenge the thesis that Sephin1’s neuroprotective properties are mediated through the prolongation of the phosphorylation of eIF2α. Instead, our results align better with the conclusions of an increasing number of recent studies indicating that Sephin1 confers neuroprotection by attenuating the IRE1α branch of the UPR^36,37^, or by blocking the NMDA-induced neuronal death^53^.

The therapeutic benefit of Sephin1 in SOD1G93A mice, in light of the negative outcome observed with guanabenz, a similar compound that additionally acts as an alpha-2-adrenergic receptor agonist, further reinforced the notion that the activation of alpha-2 adrenergic receptors led to the adverse outcome. To further investigate this, we conducted a survival efficacy study using clonidine, a specific alpha-2-adrenergic agonist. Contrary to our expectations, clonidine not only failed to accelerate the disease progression in SOD1G93A mice, but it significantly slowed disease progression and prolonged the survival duration for the female group, while having no effect on the male group. It was beyond the scope of this study to examine how clonidine specifically benefited the female mice, but it is possible that its mechanism of action involves reducing excitotoxicity by decreasing the release of glutamate, which can ultimately prevent the death of motor neurons. Previous studies have demonstrated that clonidine can indeed reduce glutamate release in mouse models of excitotoxic injury, thereby safeguarding motor neurons from damage caused by excitotoxicity^54^.

We chose to use the SOD1G93A mouse model in these studies, because it demonstrates many key features of ALS, like motor neuron death, inflammation, astroglial and microglial activation, and, relevant to this project, an upregulation of all three UPR sensors IRE1, ATF6 and PERK^55^. However, it is important to acknowledge that this is a transgene overexpression model, and therefore may not fully reflect the human disease condition. While studying the underlying disease mechanism is possible using mutated hSOD1 gene overexpression, it is crucial to note that these mice are under significant ER stress that poses a strain on normal cell functions and these effects may not be easily reversible pharmacologically.

In our study, we investigated the PERK pathway as a potential therapeutic target for ALS using a mouse-based tunicamycin assay and the SOD1G93A mouse model. Our findings suggest that targeting PERK may not be ideal for ALS treatment, as inhibiting PERK or activating it, with GSK2606414 or salubrinal respectively, did not provide therapeutic benefits. While Sephin1 did show therapeutic effects, these were likely due to PERK-independent mechanisms. The PERK pathway is a complex and dynamic process that requires a fine balance between pro-survival versus pro-apoptotic activities for optimal cellular function and survival. Achieving this balance through a single chemical entity given chronically at a fixed dose may be unrealistic. At the same time, our results suggest that other targets within the UPR, such as the IRE1α pathway and its downstream targets XBP1 and ASK1, may be better candidates for ALS therapies based on existing evidence. Additionally, we may have indirectly uncovered a new potential neuroprotective mechanism for ALS in clonidine’s activity. Though, cautious investigation of this pathway is necessary considering the potential side effects associated with the central nervous system actions of clonidine.

In summary, while the PERK pathway remains important in ALS, our study highlights the challenges of targeting it pharmacologically for therapeutic purposes. Our results suggest that exploring other targets within, and outside, the UPR may be more promising avenues for ALS treatment.

## Acknowledgements

We would like to thank Ms. Beth Levine for meticulous review and editing of the manuscript. We also want to recognize the vigilant monitoring of our SOD1G93A colony by Carlos Maya and Matt Ferola.

## Supporting information

**S1 Fig. Molecular Structures of Cited Compounds**

**S2 Fig. GSK2606414 50 mg/kg, PO, QD Survival Efficacy Effect on Body Weight.** Group average body weight over time was less well maintained in both females (by as much as 1.5 g) and in males (by as much as 0.75 g) over time in GSK2606414-treated animals compared to vehicle-treated animals. Longitudinal data analysis of body weight over time using mixed-effects maximum likelihood regression with random effects for litter and for mouse, showed that GSK2606414-treated animals weighed, on average, about 0.9 g less than controls of the same age (*p=0.023*).

**S3 Fig. GSK2606414 50 mg/kg in Standard Chow, ad libitum Survival Efficacy Effect on Body Weight.** Group average body weight over time showed that GSK2606414-treated animals maintained body weight less well than corresponding vehicle control animals. At peak body weight GSK2606414-treated animals showed an ability to maintain body weight that was about 1.75 to 2 grams lighter than control when considering group average body weight over time.

**S4 Fig. GSK2606414 18 mg/kg, PO, QD Survival Efficacy Effect on Body Weight.** Group average body weight over time showed that female GSK2606414-treated animals maintained body weight better than corresponding vehicle control animals. At peak body weight female GSK2606414-treated animals showed an ability to maintain body weight that was about 0.5 gram heavier than control when considering group average body weight over time. Males maintained body weight less well than controls and were about 1.75 g lighter than controls over a 30-d period. Overall, there was a net reduction in body weight maintenance by about 1 g when male and female effects were combined.

**S5 Fig. Salubrinal 5 mg/kg, IP, QD Survival Efficacy Effect on Body Weight.** Group average body weight over time was better maintained over time in salubrinal-treated than in vehicle-treated animals. In both male and female salubrinal-treated animals body weight was about 0.5 g heavier than corresponding control animals up until the time of first death.

**S6 Fig. Salubrinal 15 mg/kg, IP, QD Survival Efficacy Effect on Body Weight.** Group average body weight over time was less well maintained over time in male salubrinal-treated than in male vehicle-treated animals. In male salubrinal-treated animals body weight was about 0.5 g lighter than corresponding control animals. Female salubrinal-treated animals maintained body weight in a manner similar to vehicle controls. When males and females are combined, body weight was similar to control until about 65 days of age and then less well maintained thereafter.

**S7 Fig. Sephin1 4 mg/kg, IP, QOD Survival Efficacy Effect on Body Weight**. Group average body weight over time was better maintained over time in Sephin1-treated than in vehicle-treated animals from about 85 days of age (approximate time of symptom onset). Body weight was about 0.4 g heavier than in corresponding control animals. Longitudinal data analysis of body weight over time using mixed-effects maximum likelihood regression with random effects for litter, showed that sephin1-treated animals were, on average, about 0.1 g less than controls of the same age. However, this effect was not statistically significant *(p=0.86*)

**S8 Fig. Sephin1 10 mg/kg, IP, QD Survival Efficacy Effect on Body Weight.** Group average body weight over time was maintained in a manner similar to or better than vehicle controls in sephin1-treated female animals from about 70 days of age. Body weight was as much as 0.5 g heavier than in corresponding female control animals. Male sephin1-treated animals maintained body weight less well than vehicle controls through about 105 days of age, a time well after the first animal died at age 90 days. Body weight was lighter than control by as much as 0.75 g. Longitudinal data analysis of body weight over time using mixed-effects maximum likelihood regression with random effects for litter, showed that, overall, sephin1-treated animals weighed about 0.5 g less than controls of the same age. However, this effect was not statistically significant *(p=0.24*).

**S9 Fig. Clonidine 0.5 mg/kg, IP, QOD Survival Efficacy Survival Efficacy Effect on Body Weight.** Group daily average body weight over time was better maintained (by as much as 0.5 g) over time particularly after about 90 days of age in clonidine-treated female animals compared to female vehicle-treated animals. Male clonidine-treated animals maintained body weight less well (by as much as 1 g) than male vehicle controls. When both genders were combined, clonidine-treated animals maintained body weight less well than vehicle-treated animals during the period of 70 to 100 days of age. Longitudinal data analysis of body weight over time using mixed-effects maximum likelihood regression with random effects for litter and for mouse, showed that clonidine-treated animals weighed, on average, about 0.23 g more than controls of the same age, but this effect was not statistically significant (*p=0.51*).

